# Identification of essential modules regulating T cell migration to the central nervous system in multiple sclerosis

**DOI:** 10.1101/2022.06.17.496548

**Authors:** Arek Kendirli, Clara de la Rosa, Katrin F. Lämmle, Klara Eglseer, Isabel J. Bauer, Vladyslav Kavaka, Stephan Winklmeier, Christian Wichmann, Lisa-Ann Gerdes, Tania Kümpfel, Klaus Dornmair, Eduardo Beltrán, Martin Kerschensteiner, Naoto Kawakami

## Abstract

Multiple sclerosis (MS) is a neuroinflammatory disease initiated by the infiltration of autoreactive T cells into the central nervous system (CNS). Several molecules that modulate T cell CNS infiltration in MS have been identified, but how the components of cell adhesion, migration and signalling pathways interact to execute this fundamental step in MS pathogenesis is unknown. We conducted a genome-wide *in vivo* CRISPR screen in an experimental autoimmune encephalomyelitis model of MS and identified 18 essential facilitators of T cell migration that include known targets of MS therapies. Combining *in vitro* studies with *in vivo* cell transfer and multiphoton microscopy enabled us to reveal three functional modules, centred around the adhesion molecule α4-integrin, the chemokine receptor CXCR3, and the GRK2 kinase, that are required for the migration of autoreactive CD4^+^ T cells into the CNS. Single-cell analysis of T cells from patients with MS confirmed that the expression of the essential regulators correlates with the propensity of CD4^+^ T cells to reach the CNS. Taken together, our data reveal the identity and functions of key modules that govern the critical step in the induction of MS lesions.

## INTRODUCTION

Multiple sclerosis (MS) is the most common neurological disease in young adults. In MS the cascade of tissue injury is initiated when activated autoreactive T cells infiltrate the central nervous system (CNS)(1–3). The importance of this step in MS pathogenesis is well-evidenced from studies in rodent models of the disease and in humans. The capacity of CD4^+^ T cells to induce CNS inflammation has for example been demonstrated in rodent experimental autoimmune encephalomyelitis (EAE) models, in which activated T cells recognising myelin basic protein (MBP) are transferred into naïve rodents where they induce an MS-like disease(4). Experiments in such models have delineated the migratory path of encephalitogenic T cells to the CNS(5); uncovered the compartments and cellular interactions that shape the induction of CNS inflammation(6,7) and aided in the identification of adhesion molecules, such as α4-integrin, that are required for T cell migration to the CNS(6,8). Clinical data have confirmed the importance of these processes in patients with MS, showing that many of the gene loci conferring increased risk of the disease are predicted to affect CD4+ T cell activation and differentiation(9); that there is an MS-associated immune gene signature in a subset of CD4+ T cells in monozygotic twins discordant for the disease(10); that CD4+ T cells start colonizing the CNS from early stages of the disease(11); and that therapies targeting T cell migration can be effective in ameliorating MS(12).

Despite significant advances in our understanding of MS and how to treat it, most studies to date have focused on assessing and validating the roles of molecules known to be involved in T cell trafficking. However, this has left key knowledge gaps in the field and we do lack a comprehensive understanding of the molecular cues and signalling streams that are essential for T cell entry to the CNS and may thus represent alternative targets for therapy. The advent of CRISPR gene editing technology now raises the possibility of conducting comprehensive and unbiased loss-of-function screens in disease models *in vivo*: indeed genome-wide CRISPR screens have been successfully used to answer questions related to cancer initiation, propagation and therapy(13,14), as well as to reveal the mechanisms regulating critical immunological processes including T cell activation, proliferation and fate determination(15–17). To date, this powerful approach has not been applied to the question of how T cells migrate into the CNS in MS.

Here we used a rodent MS model to ask which key sets of molecules work together to determine the migration of activated CD4^+^ T cells into the CNS. By combining an unbiased genome-wide CRISPR screen with functional *in vivo* validation studies, multiphoton microscopy, and *in vitro* mechanistic experiments, we reveal three functional modules that are responsible for facilitating CD4^+^ T cell migration into the CNS. Single cell transcriptomic analysis of CD4^+^ T cells from patients with MS showed the parallel regulation of these functional modules in migrating T cell clones. Taken together our study provides a definite molecular characterization of the central step in MS pathogenesis, the entry of autoreactive T cells to the CNS.

## RESULTS

### Genome-wide CRISPR screen identifies genes essential for autoreactive T cell migration to the CNS in EAE

To study the molecular regulation of auto-reactive T cell trafficking to the CNS in MS, we used a rat EAE model, in which MS-like disease is induced by the transfer of MBP-reactive T (T_MBP_) cells(4,7,18). We first conducted a genome-wide screen to identify candidate molecules whose deletion significantly enhanced or impaired T_MBP_ cell migration into the CNS. We transduced T_MBP_ cells in a first step with the Cas9 nuclease and enhanced green fluorescent protein (EGFP), and then with blue fluorescent protein (BFP) and a genome-wide CRISPR library containing 87,690 sgRNAs targeting 21,410 genes and 396 miRNAs, as well as 800 non-targeted (NT) control sgRNAs. We included 300 x 10^6^ T cells per replicate transduced at a multiplicity of infection (MOI) < 0.3 so that statistically most T cells would contain no more than one sgRNA(19), each different sgRNA would be present in about 1000 T cells (1000x coverage) and each gene would be targeted by four different sgRNAs. After six days, these cells were injected intravenously into naïve Lewis rats. Three days later, at the time of first EAE symptoms, we isolated T cells from the blood, spleen, spinal cord meninges and parenchyma (**Figure 1A**) and used next-generation sequencing (NGS) followed by bioinformatic analysis with the MAGeCK software(20) to compare the sgRNA distribution for each gene between each of the peripheral and CNS compartments (**Figure 1B**). To the list of genes showing a differential distribution between compartments, we applied a set of selection criteria based on effect size and statistical significance (see **Methods**), leading to the identification of 1,950 candidate target genes for a subsequent validation screen. This gene list included *Itga4*, which encodes the target of the therapeutic monoclonal antibody Natalizumab, α4-integrin (**Figure 1B**). *Itga4* was one of the top most-depleted genes in all comparisons of peripheral compartments versus the CNS, validating the ability of this approach to identify clinically-relevant molecules. Furthermore, we found that many of the same gene targets appeared to regulate the entry of autoreactive T cells to both the meninges and the CNS parenchyma in EAE, as evidenced by the strong correlation of significantly differentially regulated genes between these compartments and blood and spleen (**Figure S1**).

**Figure 1.**
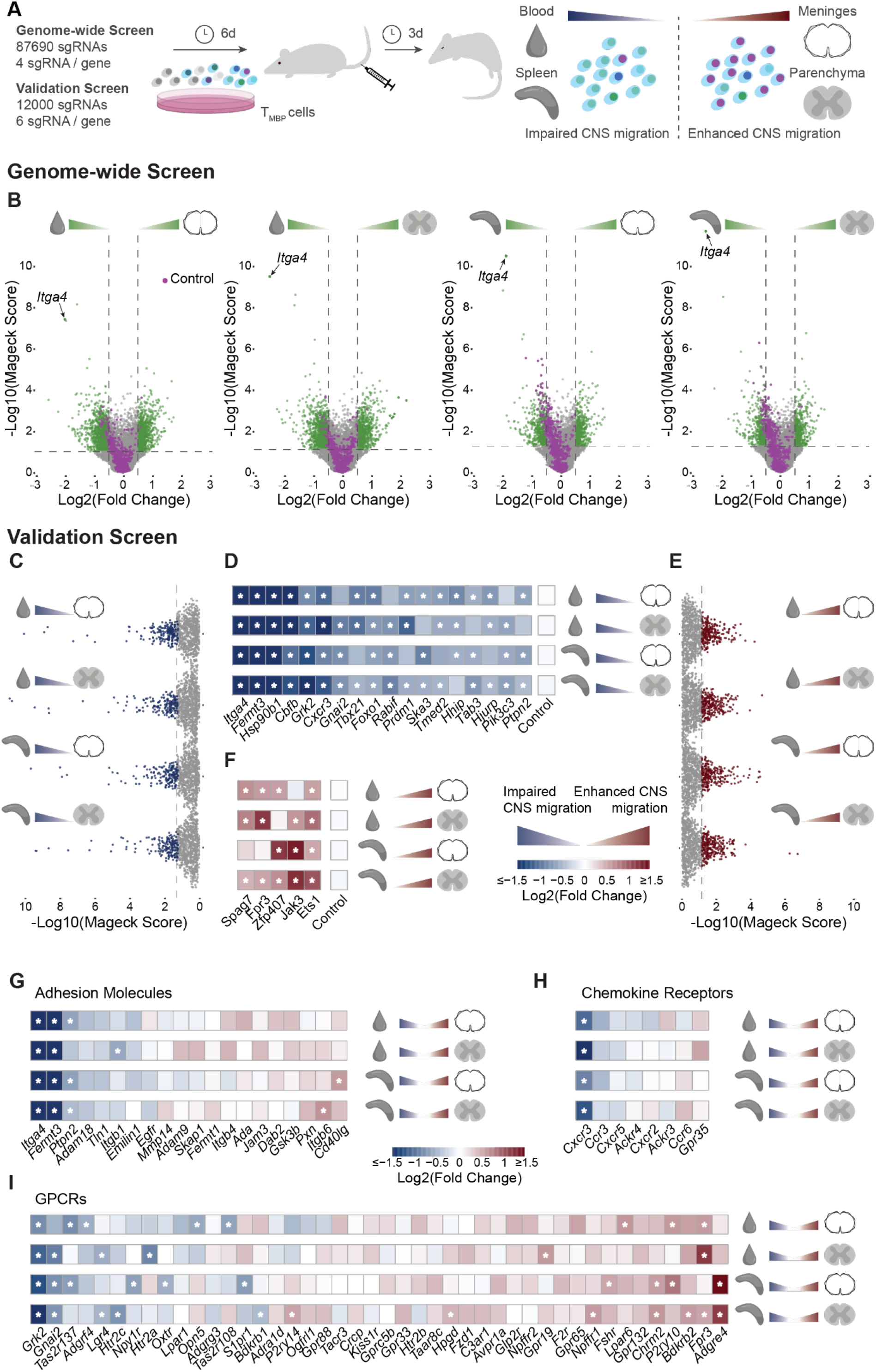
Genome-wide CRISPR screen identifies genes essential for autoreactive T cell migration to the CNS in EAE. **A**, Experimental design. T_MBP_ cells were transduced with the genome-wide or validation CRISPR libraries. After positive selection and *in vitro* reactivation, CRISPR-edited cells with single target gene-KOs were injected intravenously into Lewis rats. After three days, at disease onset, they were collected from blood, spleen, and spinal cord meninges and parenchyma for analysis. **B,** Volcano plots depicting the genome-wide screen results per gene across the four tissue comparisons. Green dots represent genes whose KO showed a sizeable change in the ability of T_MBP_ cells to migrate into the CNS. In lilac, controls. Threshold lines indicate p-value = 0.05 and log2(Fold change) = ±0.5. **C-F,** Validation screen results showing the top-ranking genes whose KO showed impaired (**C**) and enhanced (**E**) migration into the CNS, from top to bottom comparing meninges or parenchyma to blood and meninges or parenchyma to spleen, and **d, f,** log2(Fold Change) heatmaps showing the genes essential for “facilitating” T_MBP_ cell entry into the CNS (**D**) or for “braking” CNS migration (**F**). Blue indicates the gene KO showed a phenotype of impaired migration, red of enhanced migration into the CNS. Essential candidates were defined as detailed in the **Methods**. **G-I**, Log2(Fold Change) heatmaps depicting the T_MBP_ cells migratory phenotype of gene KOs as per the validation screen results for adhesion related genes (**G**) (GO terms GO.0050901, GO.0033631 and GO.0005178), chemokine receptors (**H**) (GO.0004950), and GPCRs (**I**) (GO.0004930, GO.0004703, GO.0001664 and guanine nucleotide binding genes of GO.0001664, excluding genes present in GO.0004950 or GO.0004896). For **G** and **H**, all GO term genes included in the validation screen are plotted. For **I**, only genes with a p-value < 0.01 and ≥ 3 “neg/pos|goodsgrna” as per the validation screen results are shown. Stars indicate, for all heatmaps, p-value < 0.05, absolute log2(Fold Change) > 3 standard deviations of the log2(Fold Changes) of the sample and ≥ 3 “neg/pos|goodsgrna”.

In a validation screen, we repeated the adoptive transfer experiment for the 1,950 candidate gene targets identified in the genome-wide screen, but this time targeted more stringently with six sgRNAs per gene and with enhanced selection criteria in the data analysis stage (for details see **Methods**). Based on the relative fold change of the differential distribution and its robustness across sgRNAs and different compartments, we identified 18 essential “facilitators” that are required for autoreactive T cell migration to the CNS (**Figure 1C,D**) and five essential “brakes” that limit T cell trafficking to the CNS (**Figure 1E,F**). In contrast, none of the miRNAs fulfilled these criteria, indicating that no single miRNA is essential for either promoting or preventing T cell entry to the CNS (**Figure S2**). The top ranked of the 18 essential facilitators of CD4^+^ T cell entry to the CNS belonged to the Gene Ontology (GO) terms related to “Adhesion molecules” (most prominently *Itga4* and the functionally related *Fermt3* gene, **Figure 1G**), “Chemokine receptors” (in particular *Cxcr3*, **Figure 1H**) and “G-protein coupled receptor (GPCR) related proteins” (such as *Grk2* and *Gnai2*, **Figure 1I**). Notably, some of the essential genes also encoded transcriptional regulators (**Figure S3A**), including: *Cbfb*, which forms heterodimers with RUNX proteins and has so far been primarily implicated in T cell differentiation(21); *Tbx21/T-bet*, which controls genes important for Th1 function(22); *Foxo1*, a prominent regulator of metabolic T cell fitness(23); and *Prdm1*, encoding the BLIMP1 transcription factor which is critical for both regulatory and cytotoxic T cell properties(24,25). Only one essential regulator, *Ska3*, which encodes a component of the microtubule-binding SKA1 complex(26), has a direct link to the cytoskeleton, indicating that most of the genes identified in our screen do not encode proteins that limit T cell mobility in general, but rather selectively impede trafficking from peripheral to CNS compartments.

In addition to the genes that facilitate CD4^+^ T cell entry to the CNS, our screen also detected five essential regulators, the loss of which resulted in enhanced T cell trafficking to the spinal cord meninges and parenchyma. The top ranked hit among these brakes of endogenous CNS migration was the transcriptional regulator ETS1, which regulates differentiation, survival and proliferation of lymphoid cells(27), and limits pathogenic T cell responses in atopic and autoimmune reactions(28,29). To validate the role of ETS1 in T cell migration into the CNS we used CRISPR editing to knockout (KO) the gene in T_MBP_ cells and then co-transferred *Ets1*-KO and control cells into rats and assessed their trafficking. This confirmed that *Ets1* deletion results in increased trafficking to both meninges and parenchyma (**Figure S3B,C**). Subsequent transcriptional analysis suggested that *Ets1*-deficient CD4^+^ T cells were likely to be more responsive to cytokine signals, and showed greater expression of genes encoding proinflammatory and cytotoxic mediators including IL-17 and perforin1 (**Figure S3D-I**).

As, at least in MS, the therapeutic aim is to limit T cell trafficking to the CNS, we focused our subsequent analyses on those targets that are essential for T cell entry, and in particular on the functional modules centred around key adhesion molecules, chemokine receptors and GPCR related proteins.

### The adhesion module: Itga4, Fermt3 and Hsp90b1 encode essential regulators of T cell migration to the CNS

To further characterize the “adhesion module” we first validated the effects of *Itga4*-KO on T cell migration by CRISPR editing. We co-transferred *Itga4*-KO T_MBP_ cells expressing EGFP with T_MBP_ cells edited with a NT control sgRNA and expressing BFP (control T_MBP_ cells) into rats, and collected cells from the blood and spinal cord meninges and parenchyma three days later (**Figure 2A**). By flow cytometry we confirmed that *Itga4* deletion significantly reduced T_MBP_ cell migration into the rodent CNS (**Figure 2B,C**). Accordingly, the transfer of *Itga4*-KO T_MBP_ cells alone failed to induce disease symptoms in the recipient rats, while rats that received control T_MBP_ cells showed the expected disease course (**Figure 2D**).

**Figure 2.**
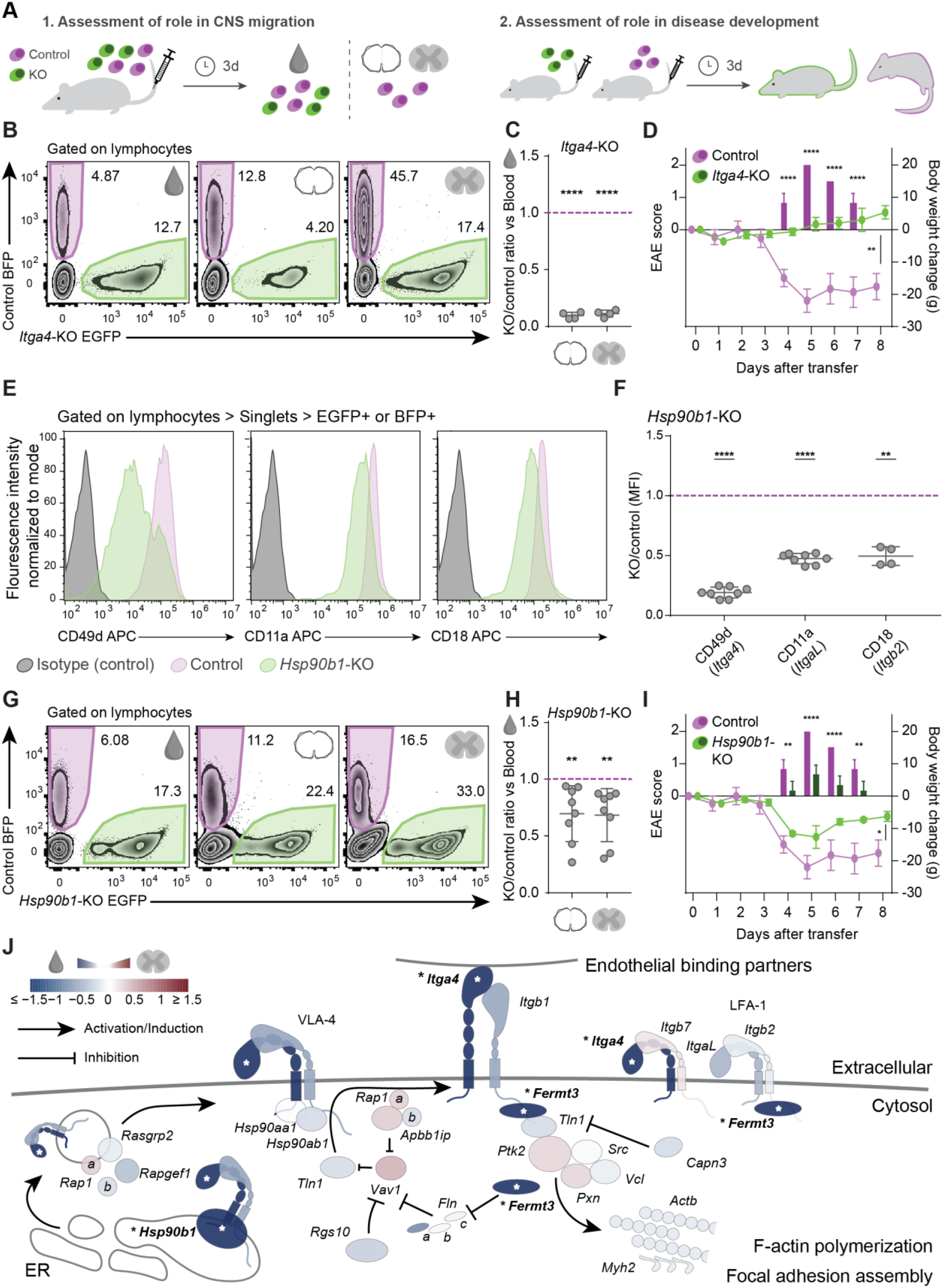
Functional validation of the adhesion module. **A,** Experimental designs. (1.) Single gene target KO-EGFP^+^ T_MBP_ cells were generated by CRISPR RNP editing, then co-transferred at a 1:1 ratio with control BFP^+^ cells into a single animal. After three days, the relative proportions of control and KO cells in blood and CNS tissues were assessed by flow cytometry. (2.) To assess the role of the gene candidates in EAE pathology, either control or single gene KO T_MBP_ cells were transferred into Lewis rats, and the disease score and body weight were evaluated over the following eight days. **B,** Representative flow cytometry plots of T_MBP_ cells recovered from blood, meninges and parenchyma after a co-transfer experiment with *Itga4*-KO and control cells. **C,** Migratory phenotype of *Itga4*-KO cells compared to control, shown as the ratio of *Itga4*-KO cell numbers/control cell numbers in meninges (left) or parenchyma (right) divided by the KO/control ratio in blood. A ratio of 1 indicates the migration behaviour of control T_MBP_ cells, a ratio below 1 indicates impaired migration into the CNS. n = 4 rats. **D,** EAE score (bars) and weight changes (line) of *Itga4*-KO-injected and control-cell-injected animals; n = 3 rats per group. **E,** Representative flow cytometry plots showing in grey the isotype-matched antibody labelling control, in lilac the control T_MBP_ cells, and in green the *Hsp90b1*-KO T_MBP_ cells, for CD49d (*Itga4*), CD11a (*ItgaL*) and CD18 (*Itgb2*). **F**, Quantification of the integrin labelling median fluorescence intensity of the *Hsp90b1*-KO cells, normalized to the control intensity. A ratio of 1 indicates the surface integrin expression levels of control T_MBP_ cells, a ratio below 1 indicates reduced integrin surface expression. n = 8 (*Hsp90b1*-KO CD49d and CD11a), n = 4 (*Hsp90b1*-KO CD18), all independent stainings. **G**, Representative flow cytometry plots of a T_MBP_ co-transfer experiment with *Hsp90b1*-KO and control cells. **H,** Migratory phenotype of *Hsp90b1*-KO cells compared to control T_MBP_ cells. n = 8 rats. **I,** EAE score (bars) and weight changes (line) of *Hsp90b1*-KO-injected and control-cell-injected animals; n = 3 rats per group. **J,** Schematic representation of the adhesion module centred around α4-integrin. The effect of each member on T_MBP_ cell migration to the CNS is color-coded based on the results of the parenchyma vs blood comparison of the CRISPR screens and stars indicate significance across at least three pairwise tissue comparisons (see **Methods**). **C, F, H,** One sample t-test or Wilcoxon signed-rank test against hypothetical mean = 1; **D, I**, repeated measures two-way ANOVA (days three to eight for disease score (**D,** F = 240.3, P = 0.0001; **I,** F = 66.13, P = 0.0012) and 0 to 8 for weight changes (**D,** F = 47.1, P = 0.0024; **I**, F = 11.53, P = 0.0274)) and Sidak’s multiple comparison test. Figures show mean ± s.d, P > 0.05 ns (non-significant), P < 0.05 *, P < 0.01 **, P < 0.001 ***, P < 0.0001 ****.

Considering other adhesion-related genes that might work alongside *Itga4* to drive T cell migration into the CNS, we next assessed the effects of knocking-out *Hsp90b1*. This heat shock protein has been proposed to act as chaperone and allow correct folding of integrins(30). Indeed *Hsp90b1*-KO T_MBP_ cells showed a marked reduction of α4-integrin surface expression and a moderate reduction of the surface expression of the LFA-1 components CD11a (integrin α-L) and CD18a (ß2-integrin) compared to control T_MBP_ cells (**Figure 2E,F)**. In contrast deletion of *Itga4* selectively abolished surface expression of α4-integrin (**Figure S4**). When *Hsp90b1*-KO T_MBP_ cells were transferred into rats, we saw a moderate yet significant reduction in trafficking to CNS meninges and parenchyma (**Figure 2G,H**), and a significantly milder EAE course (**Figure 2I**). Taken together these results outline the functional T cell adhesion module that contains α4-integrin, its intracellular binding partner kindlin3 (encoded by *Fermt3,* identified in our validation screen and with an established role in the intracellular activation of integrins(31)), and its chaperone HSP90B1 (**Fig 2J**). Notably, all other adhesion related genes in our screen are non-essential (at least by our strict definition) and even deletion of *Itgb1* that encodes ß1-integrin, which pairs with α4-integrin to form VLA-4, the binding partner for endothelial VCAM-1, shows only a mild reduction in T cell migration to the CNS, likely because it can be replaced by other ß-integrins such as ß7-integrin(32).

### The chemotaxis module: Cxcr3, Gnai2 and Tbx21 encode essential regulators of T cell migration to the CNS

CXCR3, the receptor for CXCL9, CXCL10 and CXCL11 was the only top hit in our screen among the chemokine receptor family, despite previous work implicating both CXCR3 and CCR5 as regulators of T cell trafficking in the leptomeninges(7). We first confirmed the essential role of CXCR3, showing that *Cxcr3*-KO T_MBP_ cells migrated significantly less to the CNS meninges and parenchyma compared to co-transferred control T_MBP_ cells (**Figure 3A,B**). Accordingly, transfer of *Cxcr3*-KO T_MBP_ cells induced significantly milder disease than did control T_MBP_ cells **Figure 3C**).

**Figure 3.**
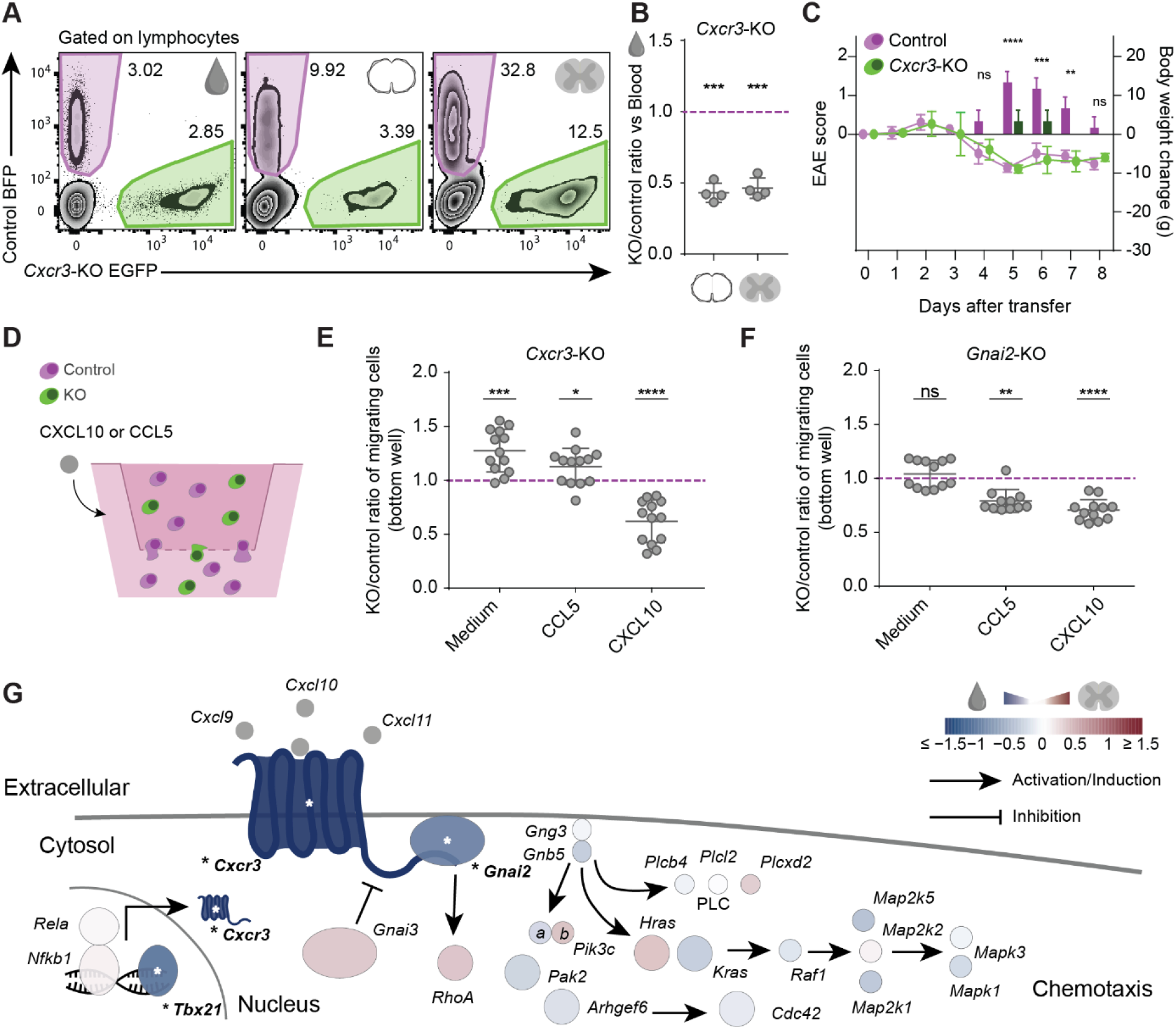
Functional validation of the chemotaxis module. **A,** Representative flow cytometry plots of a T_MBP_ co-transfer experiment with *Cxcr3*-KO and control cells. **B,** Migratory phenotype of *Cxcr3*-KO cells compared to control. n = 4 rats. **C,** EAE score (bars) and weight changes (line) of *Cxcr3*-KO-injected and control-cell-injected animals; n = 3 rats per group. **D,** Experimental design of *in vitro* transwell T_MBP_ cell migration experiments. Cells were seeded in the top chamber with medium, and control medium or chemokines were added to the bottom chamber. **E,** Migration in response to chemokine gradient of *Cxcr3*-KO T_MBP_ cells and **F**, *Gnai2*-KO cells compared to control. A ratio of 1 indicates the migration behaviour of control T_MBP_ cells, a ratio below 1 indicates impaired migration, a ratio above 1 enhanced migration. n = 13 transwell assays (*Cxcr3*-KO), n = 12 (*Gnai2*-KO medium and CXCL10), n = 11 (*Gnai2*-KO CCL5). **G**, Schematic representation of the CNS chemotaxis module centred around CXCR3 (see **Methods)**. **B, E, F,** One sample t-test or Wilcoxon signed-rank test against hypothetical mean = 1; **C**, repeated measures two-way ANOVA (days three to eight for disease score (F = 64.8, P = 0.0013) and 0 to 8 for weight changes (F = 0.1725, P = 0.6992)) and Sidak’s multiple comparison test. Figures show mean ± s.d, P > 0.05 ns (non-significant), P < 0.05 *, P < 0.01 **, P < 0.001 ***, P < 0.0001 ****.

Further analysis of this “chemotaxis module” suggests that, in addition to CXCR3, two key proteins might be involved in chemoattraction of T cells into the CNS: the guanine nucleotide binding protein GNAI2, which is linked to the intracellular transduction of signals induced by CXCL9, CXCL10 and CXCL11(33); and TBX21/T-BET, which controls the expression of the CXCR3 receptor(22). To assess the role of GNAI2 in CXCR3-mediated migration, we compared the migration of *Cxcr3*-KO and *Gnai2*-KO T_MBP_ cells to control T_MBP_ cells in a transwell chemotaxis assay. We found that *Cxcr3*-KO T cells showed altered transmigration in response to the CXCR3 ligand CXCL10, but not to the unrelated chemokine CCL5, while *Gnai2*-KO reduced migration towards both chemokines (**Figure 3D-F**). This indicates that GNAI2 transduces the effects of CXCR3 on T cell migration, while none of the other mediators of CXCR3 downstream signalling, which includes the phospholipase C (PLC), mitogen-activated protein kinase (MAPK) and RhoA pathways, is essential for the entry of autoreactive CD4^+^ T cells to the CNS in our model. Thus, based on known roles from the literature and our own experiments, the chemotaxis module comprises CXCR3, GNAI2 and the transcription factor TBX21/T-BET (**Figure 3G**).

### The egress module: GRK2 controls T cell attraction to the blood via S1PR1

Among the GPCR related proteins, the G protein coupled receptor kinase 2 (GRK2) is the dominant hit in our screen with one of strongest effect sizes of any regulator outside the adhesion module (**Figure 1D,I**). We confirmed the essential role of GRK2 by the transferring CRISPR-edited *Grk2*-KO T_MBP_ cells, which showed a significant reduction in their capacity to reach either CNS meninges or parenchyma (**Figure 4A,B**), and induced significantly milder disease symptoms (**Figure 4C**), compared to control T_MBP_ cells. As GRK2 has multiple potential target substrates and mechanisms(34), we next asked which aspect of the transmigration process was affected by loss of GRK2. We performed *in vivo* multiphoton imaging of the rat spine three days after co-injection of *Grk2*-KO T_MBP_ cells expressing EGFP and control T_MBP_ cells expressing BFP, allowing us to visualise and enumerate both types of labelled cells within the CNS (**Figure 4D**). By tracking the location and movement of individual GRK2-deficient and –competent autoreactive T cells in the meninges, we saw that GRK2 loss primarily affects the distribution of T cells between the intravascular and extravascular CNS compartments (**Figure 4E,F** and **Supplementary Movies 1** and **2**), while the speed and the path lengths of crawling T cells along the vascular surface were mostly unaffected (**Figure 4G**). This argues that loss of GRK2 alters the responsiveness of T cells to signals that determine their diapedesis but does not affect either their adhesion to the endothelial cells or their overall movement properties. These results are reminiscent of the altered trafficking of GRK2-deficient B cells between blood and lymph nodes that has been related to the desensitization of the S1PR1 receptors by GRK2(35). Therefore we next asked whether this mechanism was also important for T cell migration to the CNS using co-transfer experiments (**Figure 4H**). We found that, while *Grk2*-KO T cells showed a marked impairment in trafficking from blood to either CNS compartment, these deficits were ameliorated if T cells lacked both *Grk2* and *S1pr1* (**Figure 4I**). As trafficking from the blood to CNS was unaltered in T_MBP_ cells deficient for *S1pr1* alone (**Figure 4I)**, these findings demonstrate that the essential contribution of GRK2 to the CNS entry of T cells is mediated via S1PR1.

**Figure 4.**
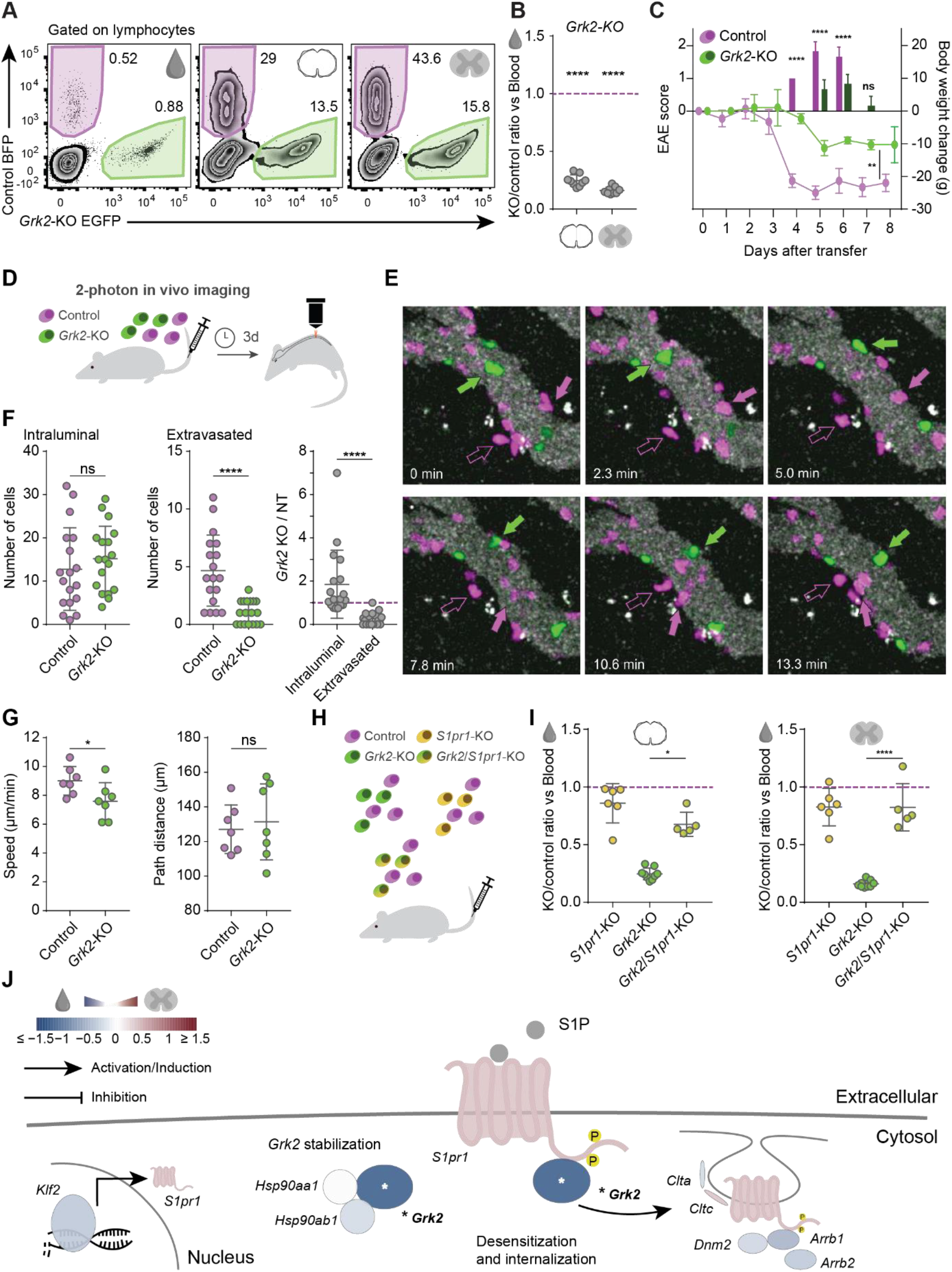
Functional validation of the egress module. **A,** Representative flow cytometry plots of T_MBP_ cells recovered from blood, meninges and parenchyma after a co-transfer experiment with *Grk2*-KO and control cells. **B,** Migratory phenotype of *Grk2*-KO cells compared to control, shown as the ratio of *Grk2*-KO cell numbers to control cell numbers in meninges (left) or parenchyma (right) divided by the KO/control ratio in blood. A ratio of 1 indicates the migration behaviour of control T_MBP_ cells, a ratio below 1 indicates impaired migration into the CNS tissue. n = 11 rats. **C,** EAE score (bars) and weight changes (line) of *Grk2*-KO-injected and control-cell-injected animals; n = 3 rats per group. **D,** Experimental design for intravital 2-photon imaging. **E,** Time-lapse images tracking cells along the meningeal vasculature. Lilac indicates control cells, green indicates *Grk2*-KO cells, filled arrows point to cells inside the blood vessel lumen, empty arrows point to extravasated cells. **F,** Distribution of cells in the leptomeninges vasculature. Left, intraluminal cell count; middle, extravasated cell count; right, ratio of KO/control cells derived from the previous two panels. N = 3 animals, 6-8 images per animal. **G,** Analysis of cell movement in the lumen of blood vessels: speed (left) and path length (right). Data summarized per movie, n = 3 animals, 2-3 movies per animal, 15-76 cells per condition per movie. **H,** Experimental design for double gene KO validation in co-transfer with control cells, and compared to co-transfer with single gene KOs. **I,** Migratory phenotype of KO cells compared to control, shown as the ratio of KO cell numbers to control cell numbers in meninges (left panel) or parenchyma (right panel) divided by the KO/control ratio in blood. n = 6 rats (*S1pr1*-KO), n = 11 rats (*Grk2*-KO, same data as in panel b), n = 5 rats (*Grk2*/*S1pr1*-KO). **J,** Schematic representation of the egress module centred around the S1PR1-GRK2 interaction. The effect of each member on T_MBP_ cell migration to the CNS is color-coded based on the results of the parenchyma vs blood comparison of the CRISPR screens and stars indicate significance across at least three pairwise tissue comparisons (see **Methods**). **B,** One sample t-test against hypothetical mean = 1; **C,** repeated measures two-way ANOVA (days three to eight for disease score (F = 28.9, P = 0.0058) and 0 to 8 for weight changes (F = 23.82, P = 0.0082)) and Sidak’s multiple comparison test; **F, G**, paired parametric t-test or Wilcoxon matched-pairs signed rank test; **i**, Kruskal-Wallis test (P < 0.0001) with Dunn’s multiple comparison test for meninges, and ordinary one way ANOVA (F = 59.89, P < 0.0001) with Turkey’s multiple comparison test for parenchyma. Figures show mean ± s.d, P > 0.05 ns (non-significant), P < 0.05 *, P < 0.01 **, P < 0.001 ***, P < 0.0001 ****.

To assess whether this GRK2-S1PR1 axis is similarly operative in human T cells, we CRISPR-edited human CD4^+^ T cells isolated from the peripheral blood of healthy donors and compared the surface expression of S1PR1 in response to exposure with its ligand S1P or its agonist fingolimod (**Figure 5A**). Our results demonstrate that human T cells lacking GRK2 showed impaired internalization of S1PR1 in response to both S1P and fingolimod exposure (**Figure 5B,C**). The findings that internalization of S1PR1 receptors was only partially impaired by the absence of GRK2 indicates that additional regulatory mechanisms are in play.

**Figure 5.**
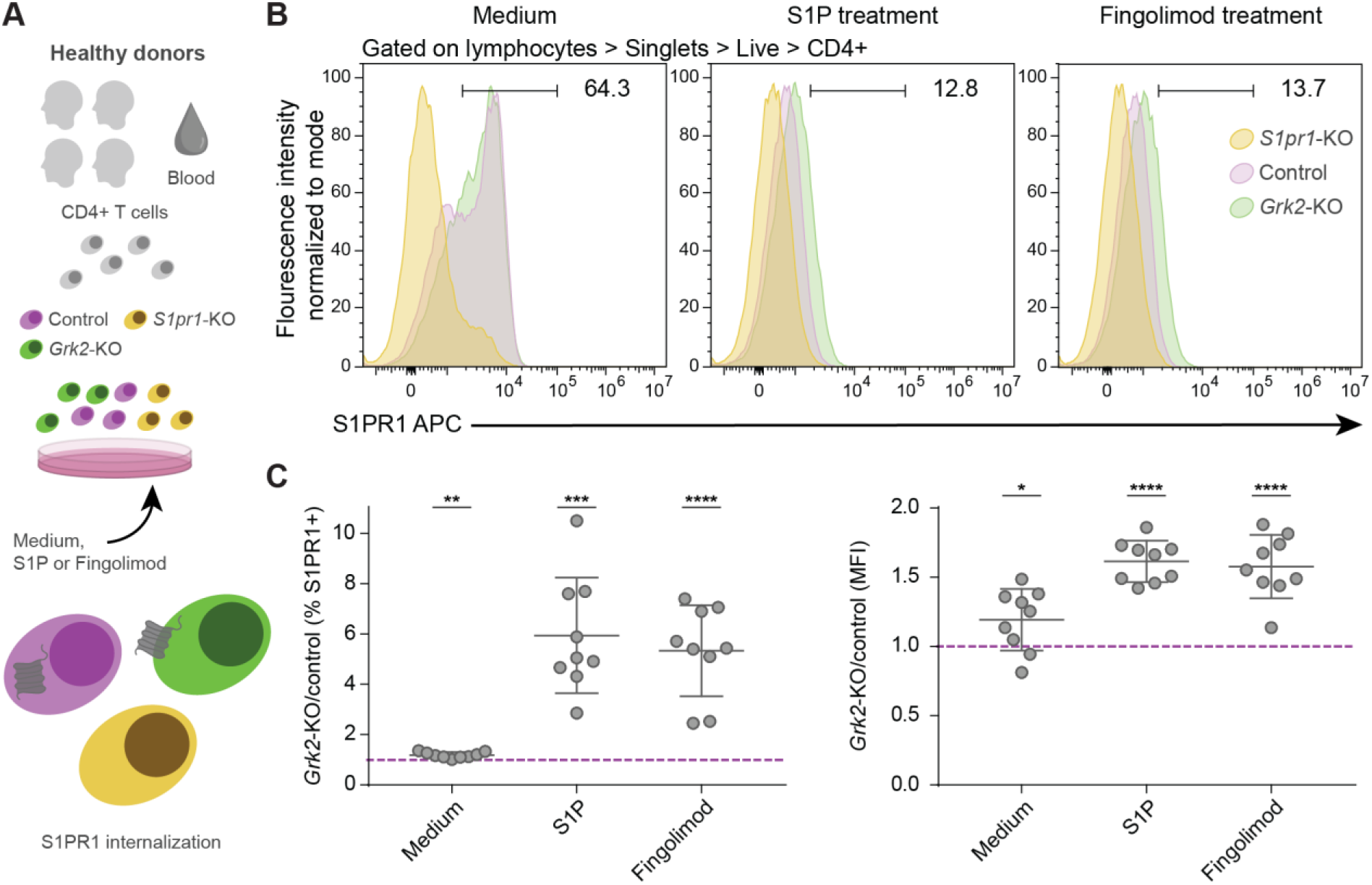
GRK2 mediates S1PR1 internalization in human T cells. **A**, Scheme of the experimental design. Briefly, CD4^+^ T cells were collected from the plasma of healthy donors and CRISPR KOs were generated *in vitro*. The cells were treated with control medium, S1P as the endogenous S1PR1 agonist, or fingolimod the MS drug, and S1PR1 internalization was measured by FACS surface staining. **B**, Representative flow cytometry plots of control and KO T_MBP_ cells for medium, S1P and fingolimod treatment conditions. **C**, Quantification of the S1PR1 internalization upon treatment. Values show the percentage of S1PR1 expressing cells (left) or the median fluorescence intensity (right) of the KO compared to the control (lilac line). A ratio of 1 indicates the surface S1PR1 expression levels of control T_MBP_ cells, a ratio below 1 indicates reduced S1PR1 surface expression, a ratio above 1 increased S1PR1 surface expression. n = 1-3 independent internalization assays from 4 donors for all groups. **C**, One sample t-test against hypothetical mean = 1. Figures show mean ± s.d, P > 0.05 ns (non-significant), P < 0.05 *, P < 0.01 **, P < 0.001 ***, P < 0.0001 ****.

Taken together, these data delineate a third functional module that controls the egress of T cells by shifting the balance of attraction between blood and CNS. This module is centred around the GRK2-mediated phosphorylation of S1PR1, which is likely induced by prolonged exposure of T cells to the S1P ligand in the blood, resulting in desensitization and subsequent internalization of S1PR1 (**Figure 4J** and **Figure 5B,C**). Our results indicate that *Grk2*-KO T cells are unable to egress from the blood even though they still receive attractive signals from the CNS, can adhere to the endothelium and can move inside blood vessels. Remarkably, the actions of S1PR1 agonists, effective therapeutic interventions in MS, also result in impaired S1PR1 signalling and reduced lesion formation in the CNS of MS patients(36). In patients treated with these agonists, however, it is assumed that the failure of T cells to respond to S1P-mediated attraction to blood sequesters these cells in peripheral lymphoid tissues before they can even reach the CNS. Here too, *S1pr1*-KO did not affect the distribution of T_MBP_ cells between blood and CNS compartments (**Figure 4I**) but led to an accumulation of these cells in the spleen (*S1pr1*-KO/control ratio in spleen vs. blood = 1.4 ± 0.25 (SD), p = 0.0291, n = 5 rats) and parathymic lymph nodes (*S1pr1*-KO/control ratio in lymph nodes vs. blood = 5.6 ± 3.24 (SD), p = 0.0336, n = 5 rats). Thus our findings reveal the delicate regulatory balance that governs S1PR1 signalling during T cell trafficking, with either sustained blocking of signal transmission (as achieved by S1PR1 agonists) or a failure to curb signal transmission (as induced by GRK2 deficiency) preventing encephalitogenic T cells from entering the CNS.

### Essential regulators identified in EAE are associated with the CNS migration propensity of T cells in patients with MS

To further assess the translational relevance of our findings we performed single-cell transcriptomic analysis of 70,594 antigen-experienced CD4^+^ T cells isolated from the blood and 16,575 such cells isolated from the cerebrospinal fluid (CSF) of four untreated MS patients and four control patients who had been diagnosed with idiopathic intracranial hypertension (IIH) (**Figure 6A**, for details see **Methods**). Bioinformatic analysis revealed twelve distinct CD4^+^ T helper cell clusters (here termed T1 to T12) that differed in their level of activation, cytotoxicity and exhaustion, as well as a cluster of regulatory T cells (Treg) defined by *FOXP3* expression (**Figure 6B,C** and **Figure S5**). All T cell clusters were present in MS patients and controls and all except for T12 were present in both blood and CSF (**Figure 6D**). To identify those T cell clusters in the blood of MS patients that are most likely to migrate to the CNS, we used their TCR sequences as tags to determine the proportion of T cell clones in each cluster that were also found in the CSF compartment of the same patient (**Figure 6E**). We then investigated whether this “CNS migration propensity” of a CD4^+^ T cell cluster correlates with the expression pattern of the functional modules of CNS migration we described above in the EAE model, and found that it did. *HSP90B1*, *GNAI2* and *S1PR1* expression level in CD4^+^ T cell clusters from patients with MS significantly correlates with their likelihood of migration into the CNS, while the expression level of *ETS1*, which limits CNS migration of T cells in rats, trends towards a negative correlation with migration propensity (**Figure 4F**). The expression of most of these regulators does not seem to shift markedly after CNS entry as their expression (with the exception of *ITGA4*, *CXCR3* and *ETS1*) was comparable in T cell clones that we present both in the blood and CNS compartment (**Figure S6**). Finally, we asked whether the expression of the functional modules of CNS migration differed between T cells isolated from the blood of MS patients and controls. We found that *FERMT3*, *HSP90B1*, *GNAI2* and *GRK2* were similarly expressed in CD4^+^ T cell clusters from patients with MS patients and controls, while the expression of *ITGA4*, *CXCR3*, *S1PR1* and *ETS1* was significantly higher in some of the T cell clusters from patients with MS (**Figure 6G**). Notably, difference in expression level of these key regulatory genes between cells from patients with MS and controls is greater in clusters with a higher migration propensity, again except for the negative regulator *ETS1* (**Figure 6H**). Taken together our single cell transcriptomic analysis of human T helper cells demonstrates that the essential regulators of T cell migration we identified in the rodent MS model are present in a sizable fraction of CD4^+^ T cells in the blood of MS patients where their expression correlates with their capacity to enter the CNS.

**Figure 6.**
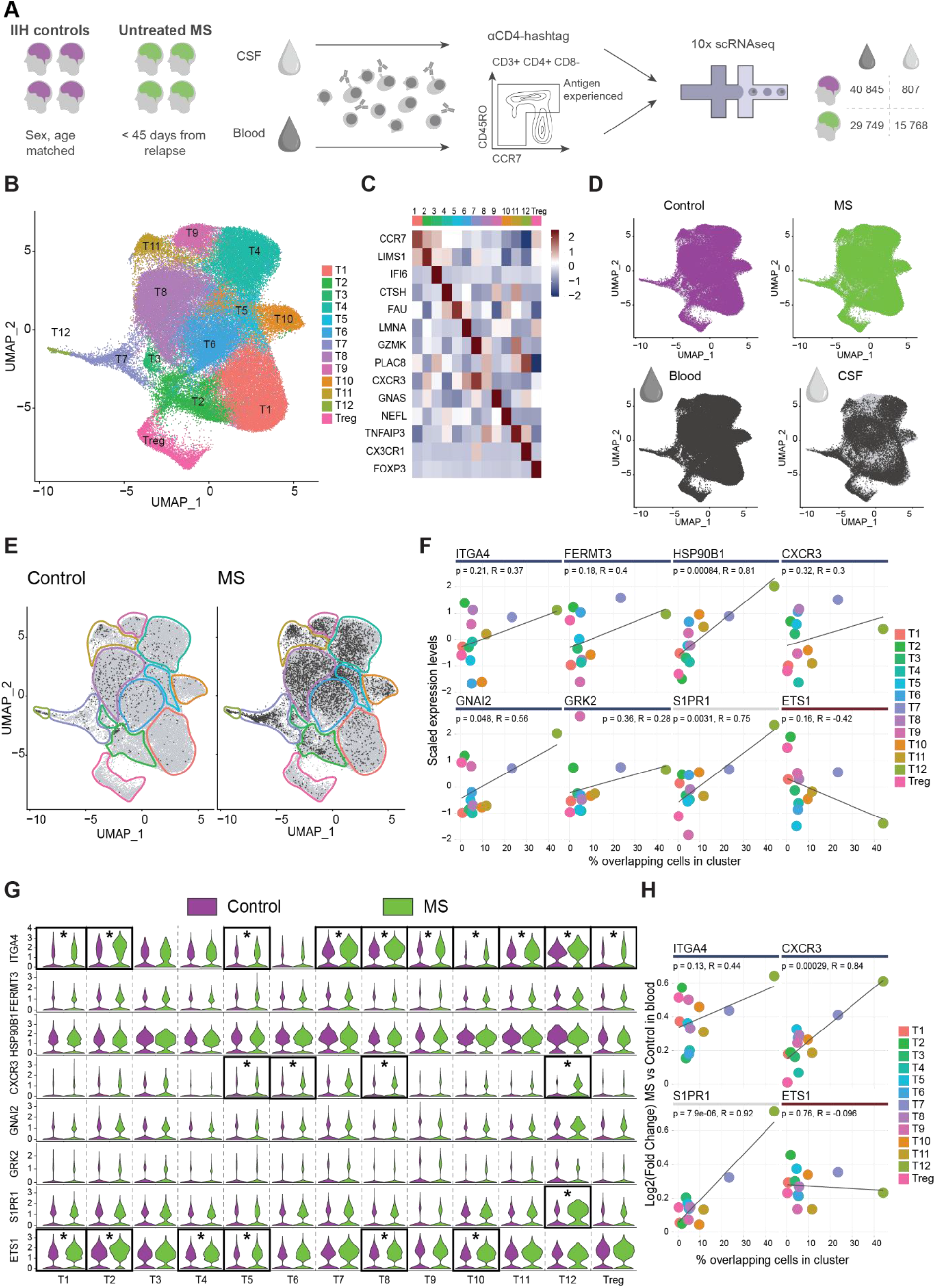
Expression of essential regulators of migration in T cells from patients with MS. **A,** scRNAseq experimental design. CD4^+^ T cells from the blood and CSF of untreated MS patients in relapse, or sex and age matched controls, were collected and analysed. **B,** CD4^+^ T cell clusters. **C,** Cluster-defining genes. **D,** Distribution of control or MS samples, and of blood or CSF samples, across the clusters. **E,** Overlapping cells in control, left, and MS, right, across clusters, defined as cells whose TCR was found in blood and CSF of the same patient. **F,** Correlations between the percentage of cells in the MS blood sample that have overlapping TCRs in the CSF, in a given cluster, and the relative gene expression level of essential regulators in this cluster in the blood of patients with MS. **G,** Violin plots of the candidate genes expression level across clusters, comparing control (lilac) and MS (green) cell samples from blood. Stars indicate significance as per adjusted p-value < 0.05 and absolute log2(Fold Change) > 3 times the standard deviation of the sample. **H,** Correlations between the percentage of cells in the MS sample that have overlapping TCRs in the CSF, in a given cluster, and the log2(Fold Change) between MS and control blood in this cluster, for those essential regulators that show differences in expression levels between MS compared to control blood as per **G**. **E, H**, R value and p-values correspond to a linear regression model.

## DISCUSSION

Here we present the results of the first genome-wide CRISPR screen aiming to identify the essential molecules that regulate the critical initial step in the formation of the MS lesion: the infiltration of autoreactive CD4^+^ T cells from the blood to the CNS. We identified three functional modules, validated the functions of the essential genes within them, and provided mechanistic evidence that brings a new level of understanding of the processes driving T cells into the disease site in both the rodent EAE model and in MS. Our data also form the foundation for future studies that extend these findings, for example by looking at the role of paracrine or systemic signals emitted by the cells in question (e.g. cytokines, chemokines or matrix metalloproteases released by the autoreactive T cells), or that search for functionally important but molecularly redundant regulatory mechanisms, neither of which can be assessed using a CRISPR screen. Beyond the study of T cell migration, the CRISPR-based screening of autoreactive T cells that we introduce here represents a highly versatile approach that can be easily adapted to interrogate the subsequent steps of T cell-mediated CNS pathology in MS, including the functional specification and survival of these cells in the target organ.

What did we learn from this unbiased and comprehensive characterization of the essential molecular signals governing autoreactive T cell entry to the CNS? First, that remarkably few molecules are *essential* for T cell migration into the CNS, and that the majority of the most potent mediators naturally cluster in three functional modules: one centred around the adhesion molecule α4-integrin, another around the chemokine receptor CXCR3, and the final one involving the GRK2-S1PR1 axis. While some of these molecules have been implicated in T cell migration in general, or even in MS, before, this is the first time that an unbiased analysis reveals all non-redundant targets among their transcriptional regulators, chaperones and binding partners as well as their intracellular signalling streams. Alongside, many previously suspected candidate regulators did not appear as “essential” in our screen. These included the chemokine receptors CCR5, CCR6 and CCR7(7,37,38), the adhesion molecules P-selectin(39), Ninjurin-1(5,40), MCAM(41), DICAM(42), the ALCAM ligand CD6(43,44) and the Lymphocyte function associated-antigen 1 (LFA1(45)). This raises interesting questions around whether these mediators are functionally redundant, or whether they are only required for a subpopulation of CD4^+^ T cells, as for LFA-1, MCAM and DICAM, which are primarily important for the migration of Th17 cells(41,42,46).

A second important finding of our screen is the presence of endogenous “brakes” of T cell migration, such as the transcription factor ETS1, which appears to limit the responsiveness of T cells to a range of immunological signals. Although these brakes are probably not therapeutic targets in CNS autoimmunity, they might be of therapeutic interest in the context of brain cancer or neurodegenerative diseases in which insufficient immune responses in the CNS can contribute to pathology(47,48).

Importantly, we showed that the essential modules we identified in the rat EAE model have high likelihood of relevance in human patients with MS. Two of the major clinical strategies that limit CNS infiltration of T cells in MS patients, blocking α4-integrin and interfering with the S1PR1 receptor, also emerge as central functional hubs in our genome-wide screen. While established S1PR1 modulators induce receptor internalization, we show here that interfering with receptor desensitization via GRK2 is an active regulatory mechanisms in human T cells and has an even more potent effect on CNS migration, at least in our rodent MS model. Interestingly, a previous study reported reduced GRK2 protein levels in PBMC isolated from relapsing-remitting and secondary progressive MS patients suggesting an altered regulation of S1PR1 internalization (49). We further show that the expression pattern of the essential regulators we identified in the EAE model reflects the propensity of defined T cell clusters from patients with MS to reach the CNS. Alongside, the expression of several of these regulators, most prominently of α4-integrin and CXCR3, is specifically higher in CNS migration-prone T cell clusters in MS patients compared to controls. These data extend and refine the results from previous population-based analyses of T cells in MS patients(50,51), and provide a molecular explanation for enhanced T cell entry to the CNS in MS. Taken together our study thus helps to define the essential molecules and modules that govern CD4^+^ T cell trafficking to the CNS and demonstrates their regulated expression in MS patients.

## AUTHOR CONTRIBUTIONS

M.K., N.K., K.F.L., C.D.R. and A.K. conceived and designed the experiments. A.K. and C.D.R. performed and evaluated CRISPR screening and editing. N.K. and K.F.L. performed and evaluated T cell transfer experiments, *in vitro* migration assays and flow cytometry analysis. N.K. and I.J.B. performed and analysed *in vivo* spinal cord imaging. L.A.G. and T.K. collected and characterized MS patient and control samples. S.W., C.W. and A.K. contributed to CRISPR editing of human T cells and K.E., V.K., K.D. and E.B performed and evaluated the single cell transcriptomic analysis. M.K. wrote the paper with input from all authors.

## ACKNOWLEDGEMENTS

We would like to thank A. Schmalz, J. Schmitt and B. Fiedler for excellent technical assistance, D. Matzek, B. Stahr, N. Ntaraklitsas for animal husbandry. We further wish to thank H. Wekerle and R. Hohlfeld for critical reading of the manuscript and L. Robinson of Insight Editing London for critical review and editing of the manuscript. We would also like to thank LAFUGA (Stefan Krebs, Helmut Blum) and CCGA Kiel (Sören Franzenburg, Janina Fuß) for NGS sequencing, the Core Facility Flow Cytometry at the BMC (Lisa Richter, Pardis Khosravani) for supporting flow cytometry and cell sorting, the Core Facility Bioimaging at the BMC (Steffen Dietzel, Andreas Thomae) for supporting the *in vivo* microscopy experiments and the LMU Biozentrum Sequencing Service for Sanger sequencing. This work was supported by a grant from the Deutsche Forschungsgemeinschaft (DFG) to N.K. and M.K. (Collaborative Research Center -TRR128, Project B10). Work in M.K.’s lab is further supported via TRR 274/1 2020 (Projects C02, C05, Z01 – ID 408885537), the Munich Cluster for Systems Neurology (SyNergy EXC 2145 – ID 390857198), the “Klaus-Faber Stiftung” and the “Verein Therapieforschung für MS-Kranke e.V.”. N.K. is further supported by a research grant from the DFG (ID 246754395) as are K.D., T.K. and E.B. (DO420/7-1). C.D.R. received support from the Fundación Rafael del Pino and is part of the DFG-funded Graduate School of Systemic Neurosciences (GSC 82 – ID 24184143).

## Contact for reagent and resource sharing

Further information and requests for resources and reagents should be directed to and will be fulfilled by the lead contact, Naoto Kawakami. Plasmids generated in this study are available upon reasonable request.

## MATERIALS AND METHODS

### Plasmids

All primer sequences are listed in **Supplementary Table 1**

The pMSCV-Cas9-EGFP vector was constructed as follows. First, a Cas9-p2a-EGFP construct was PCR amplified from the Lenti-Cas9-EGFP plasmid obtained from Addgene (63592). Then, the PCR product was assembled into an EcoRI + XhoI digested pMSCV-neo (Takara Clontech) vector by using Gibson Assembly Master Mix (NEB, E2611S). The gRNA expression vector MSCV-pU6-(BbsI)-CcdB-(BbsI)-Pgk-Puro-T2A-BFP was obtained from Addgene (86457). The pMSCV-v2-U6-(BbsI)-Pgk-Puro-T2A-GFP (with v2 improved scaffold) vector was generated as follows: first, the pU6-Pgk-Puro-T2A construct was PCR amplified from pKLV2-U6gRNA5(BbsI)-PGKpuro2ABFP-W, and the EGFP construct was PCR amplified from pMSCV-Cas9-EGFP; then, the PCR product was assembled into the SalI + XhoI digested pMSCV-neo vector using the Gibson Assembly Master Mix. The pMSCV-v2-U6-(BbsI)-Pgk-Puro-T2A-BFP (with v2 improved scaffold) vector was generated as follows: the pU6-Pgk-Puro-T2A-BFP construct was PCR amplified with overhangs for SalI + XhoI from pKLV2-U6gRNA5(BbsI)-PGKpuro2ABFP-W, then the PCR product was digested and ligated into SalI + XhoI digested pMSCV-neo vector with Quick Ligase (NEB, M2200L).

### Animals

Lewis rats were purchased from Charles River or Janvier and bred in the Core Facility Animal Models of the Biomedical Center, LMU. All animal experiments and their care were carried out in accordance with the regulations of the applicable animal welfare acts and protocols were approved by the responsible regulatory authority (Regierung von Oberbayern). All animals had free access to food and water. Animals were kept at room temperature 22 +/- 2°C, humidity 55 +/- 10 % with a Light/Dark cycle, 12h/12h (6:30-18:30). Male and female Lewis rats between 5-20 weeks old were used for the experiments.

### Generation of Cas9 expressing T_MBP_ cells

Cultures of T_MBP_ cells were established as previously described(52). Briefly, a Lewis rat was immunized in the hind-limb flanks with 100 µg guinea pig MBP (purified in-house) emulsified with complete Freund’s adjuvant (CFA) (Difco, 263810). Draining lymph nodes were collected ten days post-immunization and a single-cell suspension was prepared by passing through a metal strainer. The cells were then restimulated *ex vivo* with 10 µg/ml MBP either together with retrovirus producing GP+E86 packaging cells stably transfected with pMSCV-Cas9-EGFP, or without packaging cells, in complete DMEM supplemented with 1 % rat serum. Two days later, TCGF (complete DMEM supplemented with 10 % Horse serum and 2 % of PMA-stimulated EL4IL2 cell culture supernatant and 1 % Pen/Strep) was added to expand the number of T cells. After four days of expansion culture, the T cells were restimulated for two days with 50 Gy irradiated thymocytes in complete DMEM supplemented with 1 % rat serum and 10 µg/ml MBP, which was followed by another round of expansion in TCGF for four days. This cycle of restimulation and expansion can be repeated before experiments. In addition, successfully transduced T cells were enriched by adding 400 µg/ml neomycin for eight days from four days after first stimulation and sorting for GFP^+^ cells with BD FACSAria IIIu. The antigen specificity of the cultured T cells was confirmed by a proliferation assay, as described previously(18). Cells were frozen in a 10 % DMSO/90 % FBS mixture at -80°C, or stored in liquid nitrogen for long-term storage.

### Genome-wide rat gRNA library construction

A list of sgRNAs targeting genes and miRNAs in the rat genome was kindly provided by the Functional Genomics Consortium of The Broad Institute, Massachusetts, USA. All sgRNA sequences that were selected, and the library cloning primers, as well as protocols, are listed in **Supplementary Table 1**.

For the genome-wide library, for the vast majority of cases four sgRNA per gene or miRNA were selected, in some cases e.g. for miRNAs only fewer unique sgRNAs were available. A total of 87,690 oligos were purchased from Twist Bioscience, each as a 79-mer with a sequence of 5’- GCAGATGGCTCTTTGTCCTAGACATCGAAGACAACACCGN_20_GTTTTAGTCTTCTCG TCGCC-3’, N_20_ indicating the sgRNA sequence. Library cloning was performed as previously described(53) with minor modifications to the primer sequences and protocol (**Supplementary Table 1**): briefly, oligo pools were dissolved in TE buffer at a 10 ng/μl stock concentration, then the single-stranded oligos (1 ng) were PCR amplified for 10 cycles with Q5 High-Fidelity DNA Polymerase (NEB, M0491L) using Oligo_Amp_F and Oligo_Amp_R primers to generate double-stranded DNAs. A total of 24 reactions were pooled. The PCR products were purified with the Nucleotide Removal Kit (Qiagen, 28304). Amplified double-stranded DNAs were digested with FastDigest BpiI (BbsI: Thermo Fisher, FD1014) for 2 h at 37°C in a total of 20 reactions, and then purified with the Nucleotide Removal Kit. Ligation was performed with a T4 DNA Ligase (NEB, M0202T) using a 3 ng insert and 40 ng BpiI-digested MSCV-pU6- (BbsI)-CcdB-(BbsI)-Pgk-Puro-T2A-BFP for 16 h at 16°C per reaction in a total of 30 reactions. The ligated product was cleaned with a PCR Purification Kit (Qiagen, 28104) and the concentration was measured with Qubit 4 (Thermo Fisher). 10 ng of the ligated product was transformed into 50 μl of NEB Stable Competent cells (C3040I) in a total of 45 reactions and incubated at 30°C overnight. A Library representation above 100x was confirmed by plating transformed competent cells in serial dilutions. The plasmid DNA was prepared with an Endofree Plasmid Maxi Kit (Qiagen, 12362). For the validation library, to increase the confidence of the hits identified by the genome-wide screen, six sgRNAs were used per gene whenever possible. An oligo pool containing 12,000 oligos was purchased from Twist Bioscience and plasmid DNA was prepared similarly to the genome-wide library. For individual sgRNA cloning, two complementary oligos with BbsI-compatible overhangs were annealed and cloned into the BbsI-digested gRNA cargo plasmid by Quick Ligase. Ligated plasmids were then transformed into Stellar competent cells (Takara Clontech, 636763) and single clones were picked on the next day. The correct insertion was confirmed with Sanger sequencing (sequencing service, LMU Biozentrum) using hU6 primer.

### Gene editing of T_MBP_ cells

For viral transduction of CRISPR sgRNAs, 1.2×10^6^ HEK293T cells in complete DMEM with 10 % FBS were plated into a 10 cm diameter culture dish 18-24 h before transfection. For the transfection, 6 µg pMSCV retroviral plasmid and 3.5 µg pCL-Eco packaging vector were preincubated in 500 µl of complete DMEM at room temperature for 15 min before mixing with 500 µl of 80 µg/ml PEI max (Polysciences, 24765) in complete DMEM. After 20 min, the solution was added dropwise to HEK293T cells. The cells were cultured in 5 % CO_2_ at 37°C for 24 h and the medium was replaced with 8 ml complete DMEM + 10 % FBS for detoxification. The cells were cultured in 10 % CO_2_ at 37°C and retrovirus-containing supernatant was collected at 48 h and 72 h after transfection. The supernatant was then passed through a 0.45 µm filter to remove debris and virus was concentrated using Amicon Ultra-15 Centrifugal Filter Units (Sigma, UFC9100). For this, 14 ml supernatant was centrifuged at 4000 g for 20 min at room temperature. The concentrated virus was used immediately for transduction of rat T cells.

For transduction, freshly restimulated T cells were resuspended in DMEM with 25 mM HEPES and then enriched by Nycoprep gradient centrifugation at 800 g for 10 min at room temperature. The T cells were resuspended at concentration of 4×10^6^/ml in TCGF with 8 µg/ml polybrene (Sigma, 107689) and this suspension was plated at 500 µl/well in 12-well plates. Finally, concentrated virus solution was added at 50 µl/well and plates were centrifuged at 2000 g, room temperature for 90 min. TCGF was added at 1 ml/well and T cells were further cultured in 10 % CO_2_ at 37°C. T_MBP_ cells were transduced at a maximum multiplicity of infection (MOI) of 0.3 to prevent multiple integrations, and enough T cell numbers were kept at all times to ensure a minimum 1000x coverage (1000 T cells having the same sgRNA; 90 x 10^6^ cells for the genome-wide screen, and 12 x 10^6^ cells for the validation screen). On the next day, puromycin was added at a final concentration of 0.5 µg/ml to select for transduced T cells. After four days of culture, the T cells were restimulated as described above for two days, before injection into recipient animals. For T cell transfer, a minimum coverage of 1000x was maintained per replicate.

For candidate gene validation, CRISPR sgRNAs were introduced by RNP electroporation into previously BFP or EGFP retrovirus transduced T_MBP_ cells. sgRNAs were designed by using the GPP sgRNA designer and synthesized by Integrated DNA Technologies. The Cas9 protein and sgRNA were electroporated into the T cells by using Amaxa 4D-Nucleofector System and P4 Primary Cell 4D-Nucleofector® X Kit S (Lonza, V4XP-4032) according to manufacturer’s instructions. Briefly, to prepare the transfection reagent, 0.75 µl of Alt-R CRISPR-Cas9 tracrRNA (200 pmol/µl) and 0.75 µl of Alt-R CRISPR-Cas9 crRNA (200 pmol/µl; IDT, 1081060) were mixed; the solution was then incubated at 95°C for 5 min, decreasing to 70°C at the rate of 0.5°C/sec, at 70°C for 1 min, then cooled to 22°C. After adding 7.5 µg Alt-R S.p. HiFi Cas9 Nuclease V3 (IDT 1081058), the mixture was incubated at room temperature for 20 min. The master mix was prepared by mixing 18 µl of P4 primary solution, 4 µl of Supplement 1 and 1 µl of electroporation enhancer (stock:100 µM; IDT, 1075916). After washing with PBS 2×10^6^ T cells were pelleted and resuspended in 21 µl of master mix. This cell suspension was mixed with the transfection reagent and transfered into the Nucleofection cuvette. Electroporation was performed using the pulse code CM137.

### Tide assay

All single KO lines used in validation experiments were validated for the KO efficiency prior to the experiment (**Supplementary Table 1**). To assess the extent of genetic editing at DNA level for single sgRNAs, cells were lysed by Lysis Buffer (H_2_O with 100 µL/ml 1M Tris, 10 µL/ml 0.5 M EDTA, 40 µL/ml 3 M NaCl, 5 µL/ml Tween 20) supplemented with 5 µL/ml Proteinase K for 15 min at 56°C and 10 min at 95°C followed by cooling on ice. Then 350 µL isopropanol were added, incubated for 10 min at room temperature and centrifuged for 10 min at 16000 g at 4°C. The pellet was washed by addition of 1 ml of 70 % ethanol and 5 min centrifugation at 16000 g at 4°C. After removal of liquid the pellet was dried for 10 min at 56°C and resuspended in 50 µL water. PCR amplification of the modified DNA sequence was performed with specific primers for each target using 2x Optima PCR HotStart Polymerase (FastGene). PCR products were run on a 1 % agarose gel and amplicons of the expected size were purified using the Wizard SV Gel and PCR Clean-Up System (Promega, A9282). The samples were submitted to Sanger Sequencing (sequencing service, LMU Biozentrum). The INDELs and KO efficiency were assessed by the ICE v2 software tool(54).

### Adoptive transfer EAE and isolation of T_MBP_ cells

EAE was induced in rats by intravenous (i.v.) adoptive transfer of *in vitro* activated T_MBP_ cells. Following transfer, body weight and the EAE score of the rats were monitored daily. The EAE score was evaluated as follows: 0, no clinical signs; 0.5, partial tail weakness; 1, tail paralysis; 1.5, gait instability or impaired righting ability; 2, hind limb paresis; 2.5, hind limb paresis with dragging of one foot; 3, total hind limb paralysis. The number of transferred T cells was about 12×10^6^ for screening (a minimum of eight animals were pooled for a single genome-wide replicate and a minimum of six animals were pooled per validation screen replicate), a mixture of 3×10^6^ control T cells and 3×10^6^ KO T cells for evaluating migration into the CNS, and 1×10^6^ T cells for assessment of effects on the clinical course.

For screening and *in vivo* migration analysis, the animals were sacrificed on day three after T cell transfer, when animals showed first clinical symptoms, such as body weight loss and/or a mild clinical score (<1). Blood was drawn by heart puncture into a heparinized syringe. Spleen, parathymic lymph nodes, and leptomeninges and parenchyma of the spinal cord were dissected and homogenized by passing through a metal strainer. Lymphocytes were isolated from blood by a Nycoprep gradient. First, the blood was diluted with an equal volume of PBS and overlaid onto Nycoprep. After centrifugation at 800 g, room temperature for 30 min with mild acceleration and brake, lymphocytes were collected from the interface. For spleen, erythrocytes were removed by treating with ACK buffer for 3 min on ice and CD4^+^ T cells were enriched using the EasySep™ Rat CD4^+^ T Cell Isolation Kit (StemCell technologies, 19642), before purification by sorting. From the spinal cord parenchyma the lymphocytes were isolated using a 30 %/64 % Percoll gradient and centrifugation at 1200 g, room temperature for 30 min with mild acceleration and brake. Lymphocytes were collected from the interface.

### Flow Cytometry

For the CRISPR screens, BFP^+^ T_MBP_ cells were sorted from spleen only (genome-wide screen) or all four tissues (validation screen) using a FACS Aria III (BD) or FACS Fusion (BD) at the Flow Cytometry Core Facility of the Biomedical Centre, LMU. Cells were sorted to achieve an sgRNA coverage of >100x (the same sgRNA being present in at least 100 sorted T cells). The expression of cell surface molecules was measured by flow cytometry after antibody labelling. T cells were incubated for 30 min on ice with the 1^st^ antibody diluted in FACS buffer (PBS with 1 % rat serum and 0.05 % NaN3). After washing three times with FACS buffer, the cells were incubated with a fluorochrome-conjugated secondary antibody in FACS buffer for 30-45 min. The following antibodies were used, Mouse IgG1 Isotype control (Sigma M-1398, clone MOPC31c), mouse anti-rat CD49d (Thermo Fisher MA49D7, clone TA-2), mouse anti-rat CD11a (Biolegend 201902, clone wt.1), mouse anti-rat CD18 (Thermo Fisher MA1817, clone wt.3), APC conjugated Donkey anti-mouse IgG (Jackson ImmunoResearch 715-136-151, polyclonal). Cells were then washed once in FACS buffer and once with PBS, then resuspended in PBS and analysed by flow cytometry with Cytoflex S (Beckman Coulter). For the *in vivo* migration assay and cell surface molecule detection, BFP^+^ T_MBP_ cells and EGFP^+^ T_MBP_ cells were analysed using a FACS VERSE (BD) or LSRFortessa (BD) flow cytometer.

For *in vivo* migration assays, the KO/control ratio based on the numbers of BFP^+^ or EGFP^+^ cells in the lymphocyte gate was calculated for all tissue samples and normalized to the blood ratio of the same animal. Paired values by animal after normalization were used for statistical analysis. When plotting the fold change difference between tissues, the following transformation was applied: KO/NT ratio vs Blood = (number of KO cells in meninges or parenchyma / number of control cells in meninges or parenchyma) / (number of KO cells in blood / number of control cells in blood). Thus, the KO migration phenotype is always relative to the behaviour of the control cells within that animal, to correct for inter-animal variations.

### NGS sgRNA library preparation

Genomic DNA (gDNA) from lymphocytes or sorted T_MBP_ cells was isolated with the DNeasy Blood and Tissue Kit (Qiagen, 69504). A one-step PCR amplification was performed with Q5 High Fidelity DNA Polymerase by using 2.5 µg of gDNA per reaction with Fwd-Lib (mix of 8 staggered primers) and Rev-Lib (consists of 8 bp of unique barcode) primers for a total of 24 cycles. Illumina adapters were introduced together with the amplification primers. All primer sequences are listed in **Supplementary Table 1**. The amplified DNA amplicons were purified with SPRIselect (Beckman Coulter, B23317) with a ratio of 1:0.8 (DNA to beads) and eluted in nuclease-free water. The presence of ∼250 bp DNA amplicons was confirmed and the concentration was measured with Agilent Bioanalyzer on DNA 1000 Chips (5067–1504). Library samples were sent to The Laboratory for Functional Genome Analysis (LAFUGA) at the Gene Center Munich for sequencing single-end 50 bp on a HiSeq 1500.

### Schematic representation of functional modules

For **Figures 2h,l** and **3j**, the schematic representations of the essential modules regulating T cell migration were composed based on published studies, for the adhesion module(30,55,64–72,56–63), for the chemotaxis module(22,33,73–76), and for the egress module(35,77–81). Colour-coding corresponds to parenchyma vs blood log2(Fold Change) values derived from the validation screen for those genes included in this screen; if the gene was not included due to not passing the initial selection criteria, colour coding corresponds to the parenchyma vs blood log2(Fold Change) values derived from the genome-wide screen (**Supplementary Table 2**). For genes where more than one isoform exists, only those deemed relevant based on the literature are shown. When there were no literature reports, only the isoforms included in the validation screen and thus showing a more pronounced log2(Fold Change) in the genome-wide screen were included.

### qPCR and 3’ bulk mRNA sequencing

All primer sequences are listed in **Supplementary Table 1.**

Total RNA from cells was isolated with either a RNeasy Plus Mini (Qiagen, 74134) or a Micro (Qiagen, 74034) (for less than 100k cells) kit according to the manufacturer’s protocol. For quantitative PCR (qPCR), cDNA was synthesized using the RevertAid H Minus First Strand cDNA synthesis kit (Thermo Fisher, K1632) with 100-500 ng total RNA and Oligo (dt) primers and assays were performed on the Bio-Rad CFX Connect Real-Time PCR system using SsoAdvanced™ Universal SYBR® Green Supermix (BioRad, 1725272). The β-Actin housekeeping gene was used to normalize the variability in expression level. All qPCR reactions were run in duplicate. Results were quantified using the ΔΔCt method. For 3’ bulk mRNA sequencing, the library was prepared from total RNA using the Collibri 3’ mRNA Library Prep Kits for Illumina Systems (Thermo Fisher, A38110024). Amplification of transcripts was confirmed with Agilent Bioanalyzer on DNA 1000 Chips and sent to LAFUGA for sequencing single-end 50bp on a HiSeq 1500.

### Western blotting

T_MBP_ cells were washed twice with PBS and lysed with RIPA buffer (Thermo Fisher, 89900) including inhibitors of proteases (Sigma, 05892791001) and phosphatases (Sigma, 04906845001). Pierce™ Bovine Serum Albumin Standard Pre-Diluted Set kit (Thermo Fisher, 23208) was used to calculate the protein concentration of samples. Lysates were boiled at 95°C for 5 min in a mix with Tris-Glycine SDS sample buffer (Thermo Fisher, LC2676) and reducing agent (Thermo Fisher, NP0009). Lysates were resolved in 4-12 % Tris-Glycine gels (Thermo Fisher, XP04120) for protein separation and transferred to PVDF membrane (Millipore, IPVH09120) using Mini Gel Tank and Blot Module (Thermo Fisher, A25977, B1000). Anti-ETS1 (CST, 14069S) and β-Actin (Santa Cruz, sc-47778-HRP) primary antibodies were incubated on the membrane overnight at 4°C in 5 % BSA-TBST buffer, followed by incubation with HRP-conjugated secondary antibody (Santa Cruz, sc-2357) in case of the anti-ETS1 primary antibody at room temperature for 2 h. Blots were imaged using the LiCor Odyssey® Fc system after treatment with ECL Western Blotting-Substrat (Thermo Fisher, 32209).

### Transwell chemotaxis assay

Chemotaxis assays were performed using a 96-well transwell chamber with 5 μm pore size (Corning, 3387). T cells were resuspended in complete DMEM supplemented with 1% rat serum. Control-BFP^+^ and KO-EGFP^+^ cells were counted and mixed at a ratio of 1:1. Each upper insert received 0.2 × 10^6^ T cells in 75 μL medium. To the lower compartment, 235 μL of complete DMEM supplemented with 1 % rat serum with or without chemotactic stimuli 30 ng/mL CXCL10 (PeproTech, 400-33) and CCL5 (PeproTech, 400-13) were added. The chemotaxis plates were centrifuged (400 g, 1 min) and incubated at 37°C with 10 % CO_2_ for 5 h. After incubation, migrated cells in the lower chamber were analysed by flow cytometry using LSRFortessa (BD) or CytoFlex S (Beckman Coulter). For analysis and quantification, the percentage of cells detected in the lower chamber was normalized to input values. Then, the KO/control ratio was calculated for all conditions.

### Intravital imaging of the spinal cord leptomeninges

Two or three days after intravenous co-transfer of 1×10^6^ BFP^+^ control and 1×10^6^ EGFP^+^ *Grk2*- KO T cells, intravital imaging within the spinal cord leptomeninges was performed as previously described(6). The animal was anesthetized by intramuscular injection of MMF (2 mg/kg Midazolam, 150 µg/kg Medetomidine and 5 µg/kg Fentanyl), and a tracheotomy was performed to allow mechanical ventilation with 1.5-2.0 % isoflurane in air. The body temperature of the animal was maintained by a heat-pad placed underneath. Furthermore, a catheter was inserted into the tail vein to allow intravenous injection of Texas-Red conjugated 70 kDa dextran (100 µg) to visualize blood plasma during the imaging. To allow imaging, a laminectomy was performed at the dorsal part of Th12/13. For this, the skin was opened with a midline incision of 3 cm and the paravertebral musculature on the spine was removed. Then the animal was fixed in a custom-made fixation device, which provides stability by pushing with three pins from one side to the spine. The dorsal part of the central spine disc was removed after cutting both sides using a dental drill. The dura was then removed. To avoid artifacts due to breathing, the animal was slightly lifted before starting imaging. Time-lapse images were acquired with a Leica SP8 microscope using a water-immersion 25x objective lens (N.A.: 1.00, WD: 2.6 mm). For excitation of BFP and EGFP, pulsed-laser from an InSight DS+ Single (Spectra Physics) was adjusted to 840 nm and fluorescence signals were first separated with a beam splitter BS560. Signals of shorter wavelength were again split by BS505 and detected after the band pass filters HC405/150 (BFP) and ET525/50 (EGFP). Signals of a longer wavelength were again separated by beam splitter RSR620 and detected after the band pass filter BP585/40 (Texas Red). Images were acquired from a field of approximately 440 µm x 440 µm with a resolution of 512×512 pixels and an approximate 100 µm z-stack, with an interval of 2-3 µm.

The images were processed by Fiji. First, a Gaussian Blur filter (cut off of 1 pixel) was used, followed by maximum Z-projection. When necessary, bleed-through liner subtraction was applied. Finally, signal intensity was adjusted by linearly adjusting brightness and contrast. For tracking of the cells, the Manual Tracking Plugin of Fiji was used to obtain coordinates, which were used to calculate speed and length of the cells in Excel together with information from imaging such as time and pixel resolution. Locations of cells were analysed by the Cell Counter Plugin of Fiji.

### Study patients

CSF sampling was performed to confirm the diagnosis of relapsing-remitting MS according to the revised McDonald criteria, all patients had encountered an MS relapse in the 45 days prior to lumbar puncture and had not received any disease modifying treatments (but one patient was treated with high dose steroids 24 days before sampling). CSF and blood samples from four sex- and age-matched individuals diagnosed with idiopathic intracranial hypertension were included in the control group. Patient samples were collected at the Institute of Clinical Neuroimmunology at the LMU Klinikum Munich, Germany. Recruitment of individuals took place from August 2020 to January 2021. Collection of blood and CSF was approved by the local ethics committees of the LMU, Munich (ethical vote: 163-16). Written informed consent was obtained from all subjects according to the Declaration of Helsinki. PBMCs of four healthy donors used for CRISPR gene editing were derived from leukoreduction system chambers provided by the Department of Transfusion Medicine at the LMU Klinikum Munich, Germany (ethical vote: LMU #18-821).

### S1PR1 internalization assay with human CD4^+^ T cells

Cells derived from leukoreduction system chambers were diluted 1:5 with PBS and added to a SepMate tube containing human Pancoll. During centrifugation (1200 g for 10 min at 4°C) PBMCs were isolated by density gradient. CD4^+^ Human T cells were enriched from PBMCs using EasySep™ Human CD4^+^ T Cell Isolation Kit (StemCell Technologies) according to the manufacturers protocol. CRISPR RNPs targeting control (NT), *S1PR1* and *GRK2*, were delivered into the CD4^+^ Human T cells by using Amaxa 4D-Nucleofector System and P2 Primary Cell 4D-Nucleofector® X Kit S (Lonza) according to manufacturer’s instructions using the pulse code EH100. Briefly, per reaction, 0.375 µl of Alt-R CRISPR-Cas9 tracrRNA (200 pmol/µl) and 0.375 µl of Alt-R CRISPR-Cas9 crRNA (200 pmol/µl) were mixed and the solution was then incubated at 95°C for 5 min, decreasing to 70°C at the rate of 0.5°C/sec, at 70°C for 1 min, then allowed to cool to room temperature. 5 µg Alt-R S.p. HiFi Cas9 Nuclease V3 (IDT) was mixed with crRNA:tracrRNA duplex and the mixture was incubated at room temperature for 20 min for RNP formation. After washing with PBS 2×10^6^ T cells were pelleted and resuspended in 23 µl of P2 nucleofection buffer and mixed with RNP complex. 0.8 µl of Alt-R® Cas9 Electroporation Enhancer (stock:100 µM) (IDT) was added per reaction before the electroporation. RNP electroporated T cells transferred to round bottom 96-well plate (Corning) at a concentration of 1×10^6^ T cells per well in RPMI medium supplemented with Primocin (100 µg/ml; Invivogen), 10 % charcoal stripped FBS (Thermo Fisher), monoclonal anti-CD3 antibody (1 µg/mL; Thermo Fisher) and anti-CD28 antibody (2 µg/mL; Thermo Fisher) for T cell activation for 72 h at 37°C incubator. We used charcoal stripped FBS to prevent unwanted S1PR1 receptor internalization. T cells then were washed and further incubated with recombinant human IL-2 (10 ng/ml; R&D systems) for up to 2 weeks at 37°C incubator with medium change every 3 days. Human T cells between day 6 and day 13 after isolation were then used for the S1PR1 internalization assay. T cells were incubated in medium consisting of either 1 µM S1P (Tocris) or 1 nM fingolimod-P (FTY720 Phosphate, Biomol) for 90 min at 37°C. PBS containing 4 % Fatty-acid free BSA (Sigma) which was used to dissolve S1P was added as vehicle for untreated control. T cells, then, were washed with PBS and stained with anti-CD4 (Biolegend) and anti-S1PR1 (Thermo Fisher) at a concentration of 1:100 and with LIVE/DEAD Violet dye (Thermo Fisher) at 1:1000 concentration for 30 min in FACS buffer (PBS with 1 mM EDTA). S1PR1 expression in CD4^+^ T cells was assessed by flow cytometry using CytoFlex S (Beckman Coulter).

### Collection and processing of human samples for single cell transcriptomics

#### PBMC

After collection of blood into tubes containing EDTA, samples were diluted 1:1 with PBS and added to a SepMate tube containing human Pancoll. During centrifugation (1200 g for 10 min at 4°C) PBMCs were isolated by density gradient. The isolated cells in plasma were transferred to a tube and centrifuged again at 300 g 10 min at 4°C. The isolated PBMCs were either used freshly or cryopreserved in liquid nitrogen, using serum-free cryopreservation medium (CTL-Cryo ABC Media Kit, Immunospot).

For analysis, samples were thawed quickly, washed twice with 1 % BSA/PBS (300 g for 10 min at 4°C) and labelled using the following procedure: 10 min at 4°C with Fc-block (Miltenyi #130-059-901) at 1/50 dilution in FACS buffer (PBS + 2% FBS), followed by the surface antibody mix. The mix comprised: ThermoFisher anti-human CD45RO-FITC (#11-0457-42, dilution 1:40), CCR7-APC (#17-1979-42, 1:40), CD3-AF700 (#56-0037-42, 1:50), Fixable Viability Dye eFluor™ 780 (#65-0865-14, 1:1000) at 1:1000, CD4-Pacific Blue (#MHCD0428, 1:25) and BioLegend: CD8-PerCP (#344708), in a total volume of 100 µl FACS buffer and incubated for 30 min at 4°C. Antigen-experienced CD4^+^ T cells consisting of CD3^+^CD4^+^CD8^-^ effector memory (CD45RO^+^CCR7^-^), effector (CD45RO^-^CCR7^-^) and central memory (CD45RO^+^CCR7^+^) cells were collected using a FACS Aria Fusion flow cytometer (BD Biosciences). After, cells were washed in 0.04 % BSA/PBS and approximately 16,500 cells per sample were loaded onto the 10x Chip.

#### Cerebrospinal Fluid (CSF)

Human CSF samples (3-6 ml) were processed within 1 h of lumbar puncture. After centrifugation at 300 g for 10 min, the cell pellet was incubated in a 2 ml tube with TotalSeq-C Antibodies: anti-human CD4, CD8A and mouse IgG1 isotype control (Biolegend, #300567, #301071, #400187; 0.5 µg of each) Therefore, we followed the Cell Surface Labelling Protocol from 10x Genomics, but with all centrifugations done at 300 g for 10 min. All cells were loaded on the 10x Chip, with a maximum target cell number of 10,000.

#### 10x library preparation and sequencing

Further processing followed the manufacturer’s protocol using the Chromium Next GEM Single Cell VDJ v1.1. For CSF samples, the Feature Barcoding technology for Cell Surface Protein steps were also performed. Libraries were sequenced on an Illumina NovaSeq6000 S4 using read lengths of 150 bp read 1, 8 bp i7 index, 150 bp read 2.

### Bioinformatic analysis

#### CRISPR screen analysis

The Galaxy platform(82) was used for data analysis. For raw fastq files, Je-Demultiplex-Illu was used for de-multiplexing, followed by Cutadapt and Trimmomatic to get the 20 bp sgRNA sequence. Counts were then obtained with MAGeCK(20) (version 0.5.7.1+) count. Normalization across samples was conducted in R(83) (version 4.0.0+) after a 50 raw count threshold, and sgRNAs with fewer than 50 counts in more than two replicates of the same tissue were discarded altogether. The MAGeCK test was run without normalization or zero removal, and otherwise default parameters on Galaxy, giving the information of NT sgRNA controls for noise correction. All further data processing was done with R.

#### Genome-wide screen analysis and validation screen design

For plotting of the NT values in genome-wide analysis results (**Figure 1B-E**), four random NT sgRNAs were sampled with replacement from the NT sgRNA pool, for a total of 800 combinations of NT “genes” with four different sgRNAs each, to model the behaviour of a wild-type gene based on individual sgRNAs.

The selection of candidates for the validation screen was done based on the MAGeCK results of the genome-wide screen. All genes in meninges vs blood and parenchyma vs blood comparisons with an absolute “neg|lfc” or “pos|lfc” > 0.5 were included. Genes from other comparisons (meninges or parenchyma vs spleen and spleen vs blood) were included when the absolute “neg|lfc” or “pos|lfc” > 1 and the number of “neg|goodsgrna” for negative “neg|lfc” candidates or of “pos|goodsgrna” for positive “pos|lfc” candidates ≥ 2, or the absolute “neg|lfc” or “pos|lfc” > 0.6 and the number of “neg|goodsgrna” or “pos|goodsgrna” > 2. Only genes expressed in T cells as per(7) were included for validation.

#### Identification of essential regulators of migration

Validation screen candidates were considered essential regulators of T cell migration to the CNS when they met the following criteria as per the results of the MAGeCK analysis: “neg|lfc” < -3 x standard deviation of all “neg|lfc” and “pos|lfc” in the sample, the number of “neg|goodsgrna” ≥ 3, and the “neg|p-value” < 0.05; or “pos|lfc” > 3 x standard deviation of all “neg|lfc” and “pos|lfc” in the sample, the number of “pos|goodsgrna” ≥ 3, and the “pos|p-value” < 0.05.

#### Bulk RNA sequencing data

Galaxy and R were also used for bulk RNAseq data processing. Fastq files were aligned with RNA STAR (version 2.7.2b) to the reference genome Rnor_6.0.102 with default parameters and without trimming. HTSeq-count (version 1.0.0) was then used, and differential expression determined with DESeq2 (version 2.11.40.7+galaxy1), with estimateSizeFactors = poscounts and otherwise default parameters. Batch correction per animal was performed when applicable. All further analysis was run with R(83,84) (version 4.0.0+). For pathway analysis, the gProfiler(85) web tool was used on genes with an adjusted p-value < 0.05 in an ordered query from most to least extreme log2(Fold Change).

#### Processing of single cell sequencing data

Sequencing results were demultiplexed and aligned to the human GRCh38 reference genome using Cell ranger (10X Genomics, v.6.1). Gene barcodes with unique molecular identifier (UMI) counts that reached the threshold for cell detection were included in subsequent data analysis, using the R package Seurat(86) (version 4.1.0).

#### Analysis within compartments

In a first step, data from the different compartments were analysed separately. Data from cells with between 200 and 5000 genes per cell and a percentage of mitochondrial genes below 10 % were included in further analyses. Genes expressed in fewer than three cells were excluded. Data were log-normalized and a batch effect was found in both the CSF and blood compartments. Therefore, established integration methods implemented within the Seurat package (CCA and rPCA) were applied, using the vst algorithm for detection of variable features. Data were then scaled and principal components computed for dimensional reduction. Applying k nearest neighbours, the neighbourhood overlap between cells was computed, followed by clustering the cells with a shared nearest-neighbour-based clustering algorithm. Uniform manifold approximation and projection (UMAP) was applied to visualize data in a two-dimensional space. The information from T cell receptor (TCR) enrichment library sequencing (10x Genomics, v.6.1) was added for each cell, followed by subsetting for cells where a TCR clonotype and only one beta chain had been detected. Small cell clusters with low quality and doublets, as well as a cluster expressing NKT cell signatures, were removed.

#### Computing the TCR overlap between tissues

For each individual MS or control sample, the amino acid sequence information from the TCR enrichment sequencing was used to match cells with identical TCR expression present in both the blood and CSF compartment of the same patient. If the expression of a TCR was found across compartments, cells were considered as overlapping. For TCRs found more or equal three times in total, cells were labelled as expanded across tissues.

#### Combined analysis

Both data sets were merged, log-normalised, integrated with rPCA, and scaled, before undergoing principal component analysis, neighbourhood computation, clustering and dimensionality reduction using UMAP. The remaining twelve clusters were assigned a number, based on their expression level of CCR7 from high to low. Each of the clusters expressed specific features, making them distinct on a transcriptomic level (Extended Data Figure 7). Cluster T_12 included only 13 cells from the CSF, and was manually assigned to the nearest cluster T_7 as it appeared to be unique for the blood compartment.

### Statistical analysis and software

FlowJO (version 10+) was used for analysis of flow cytometry data. For statistical analyses and plotting, GraphPad Prism version 7+ (GraphPad Software) and R(83,84) (version 4.0.0+) were used. Calculations for total cell numbers were performed using R and Excel (Microsoft Office). Unless otherwise noted, data are represented as mean ± s.d (standard deviation), sample sizes reported in the figure legends, all replicates are biological, and measurements were not repeated. KO/control comparisons are represented as a ratio KO phenotype/ control phenotype, unless otherwise specified. The statistical significances and tests are reported in the Figure legends. The Shapiro-Wilk normality test was used to determine Gaussian distribution. For Gaussian distributions of the data, parametric tests were applied, for non Gaussian, non-parametric tests. CRISPR screen result statistical analysis was performed solely with the MAGeCK software(20), and RNAseq statistical analysis was performed solely by DESeq2 (galaxy), as described elsewhere. Whenever applicable, paired statistical tests were run, as indicated in the figure legends, pairing KO and control cells of the same sample. Test statistics were corrected for multiple testing if more than three comparisons were run in parallel, with two-stage linear step-up procedure of Benjamin, Krieger and Yekutieli (fdr 1 %). For the evaluation of KO T_MBP_ cell migration validation experiments by FACS, animals with < 100 cells detected in any population were excluded. All KO/control phenotype datasets were statistically evaluated with one sample t-tests (parametric) or Wilcoxon signed-rank tests (non-parametric). For the *Grk2*/*S1pr1*-KO experiment, an ordinary one-way ANOVA with Tukey’s multiple comparison tests (parametric) and Kruskal-Wallis test with Dunn’s multiple comparison test (non-parametric) were run for the KO/control phenotypes across the different KOs. For the assessment of the disease induction and weight changes phenotypes of the KO cell transfer, a repeated measures two-way ANOVA for time and genotype variations (days three to eight after T_MBP_ cell transfer for disease score, and days zero to eight for weight changes) and Sidak’s multiple comparison test were run. Two-way ANOVA F and P-values reported in the figure legends correspond to T_MBP_ cell genotype variation (one degree of freedom), KO T_MBP_ cells compared to control T_MBP_ cells. All statistical tests and p-values are two-tailed. For the human data, differential expression of genes between MS and Control and blood and CSF was computed using the R package MAST90 within the FindMarkers function of Seurat. Correlations are either Pearson correlations or linear regression models, as indicated in the Figure legends. Both CRISPR screens have three replicates. One blood replicate from the genome-wide screen was excluded due to bad technical quality. The bulk RNAseq for the *Ets1*-KO has three replicates. All other n values are reported in the figure legends. Significance was set as P < 0.05 *. P > 0.05 ns (non-significant), P < 0.05 *, P < 0.01 **, P < 0.001 ***, P < 0.0001 ****. Adobe Illustrator (Adobe Systems), Inkscape and PowerPoint (Microsoft Office) were used for figure preparation.

### Data and code availability

Next-generation sequencing raw data and processed gene expression data that support the findings of this study will be deposited into GEO and will be made publicly available following peer review. All other data generated or analyzed during this study is included in the published article or are available from the corresponding author upon reasonable request. No custom algorithms were used for the analysis of the data for this paper. Code will be provided upon request.

## SUPPLEMENTAL INFORMATION

**Figure S1-S6**

**Movie S1. Time lapse movie on day two after TMBP cell transfer. Related to Figure 4**

Time lapse movie of control (lilac) and *Grk2*-KO TMBP cells (green) at the spinal cord leptomeninges at before onset of clinical EAE on day two after T_MBP_ cell transfer. Blood vessels (grey) were visualized by intravenous injection of fluorescent dextran. Open arrow heads indicate the extravasated control and *Grk2*-KO T_MBP_ cells. Inserted number indicates the time after start imaging. Scale bar: 50 μm.

**Movie S2. Time lapse movie on day three after TMBP cell transfer Related to Figure 4**

Time lapse movie of control (lilac) and *Grk2*-KO T_MBP_ cells (green) at the spinal cord leptomeninges after onset of clinical EAE on day three after T_MBP_ cell transfer. Blood vessels (grey) were visualized by intravenous injection of fluorescent dextran. White dashed lines outline blood vessels. Inserted number indicates the time after start imaging. Scale bar: 50 μm.

**Table S1.** Primers + Oligos + PCR protocols

**Table S2.** MAGeCK data for module schemes Related to **Figure 2J, 3G**, **4J**

ModuleSchemesMageckData

