## Supplementary Figures for "Identification of essential modules regulating T cell migration to the central nervous system in multiple sclerosis"

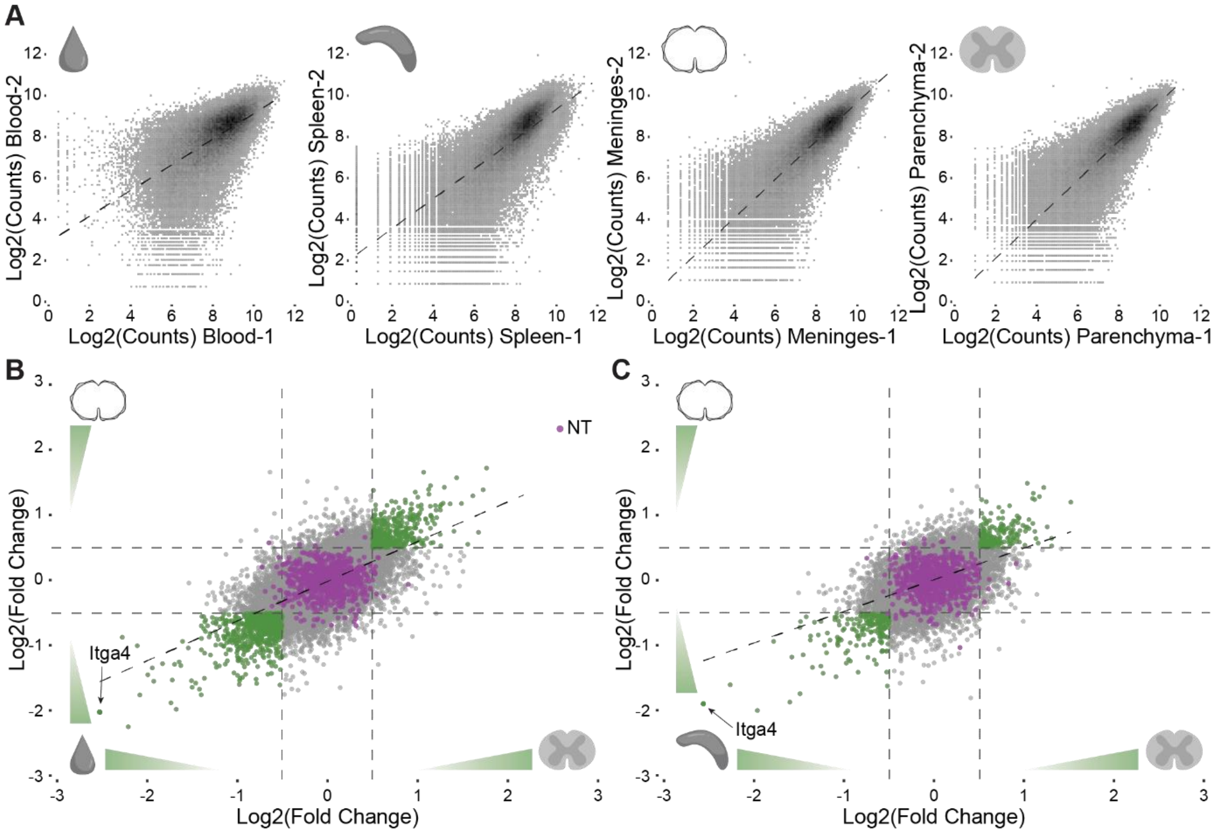


**Figure S1. Correlation analysis of genome-wide CRISPR screen**

**A,** Correlations of the raw sgRNA count values of two representative replicates in (left to right) blood (Pearson correlation index of 0.74), spleen (0.78), meninges (0.86) and parenchyma (0.85). **B,** Correlation of log2(Fold Change) of genome-wide CRISPR screen for meninges vs blood compared to parenchyma vs blood (Pearson correlation index 0.62) and **C** meninges vs spleen compared to parenchyma vs spleen (0.48). Green, correlating genes whose KO shows a sizeable change in the ability of $T_{MBP}$ cells to migrate into the CNS. In lilac, controls.

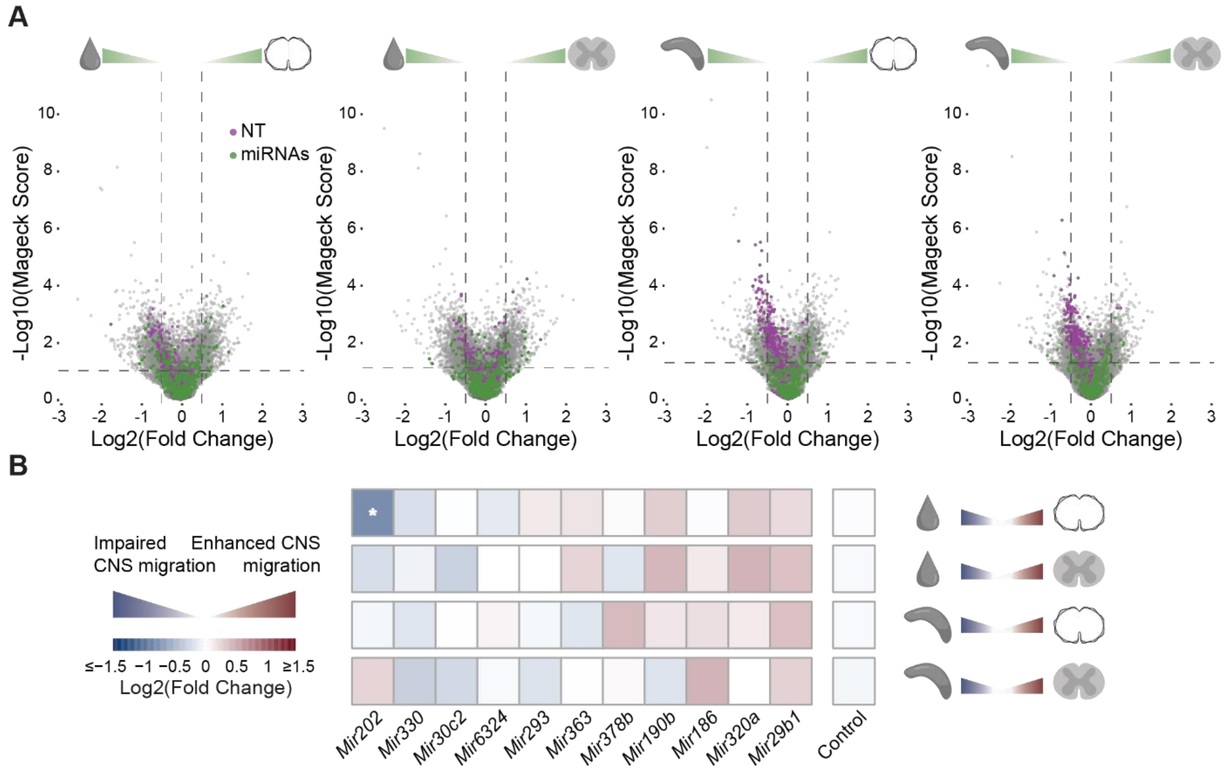


**Figure S2. miRNA based regulation of T cell migration into the CNS**

**A,** Genome-wide screen results across different tissue comparisons, comparing (from left to right) meninges or parenchyma to blood and meninges or parenchyma to spleen. Threshold lines indicate p-value = 0.05 and log2(Fold Change) = ±0.5. In green, miRNAs, in lilac, NT controls.

**B,** Results from the validation screen showing the miRNAs that were included, comparing (from top to bottom) meninges or parenchyma to blood and meninges or parenchyma to spleen. Star indicates p-value < 0.05, absolute log2(Fold Change) > 3 standard deviations of the sample and ≥ 3 "neg/pos|goodsgrna".

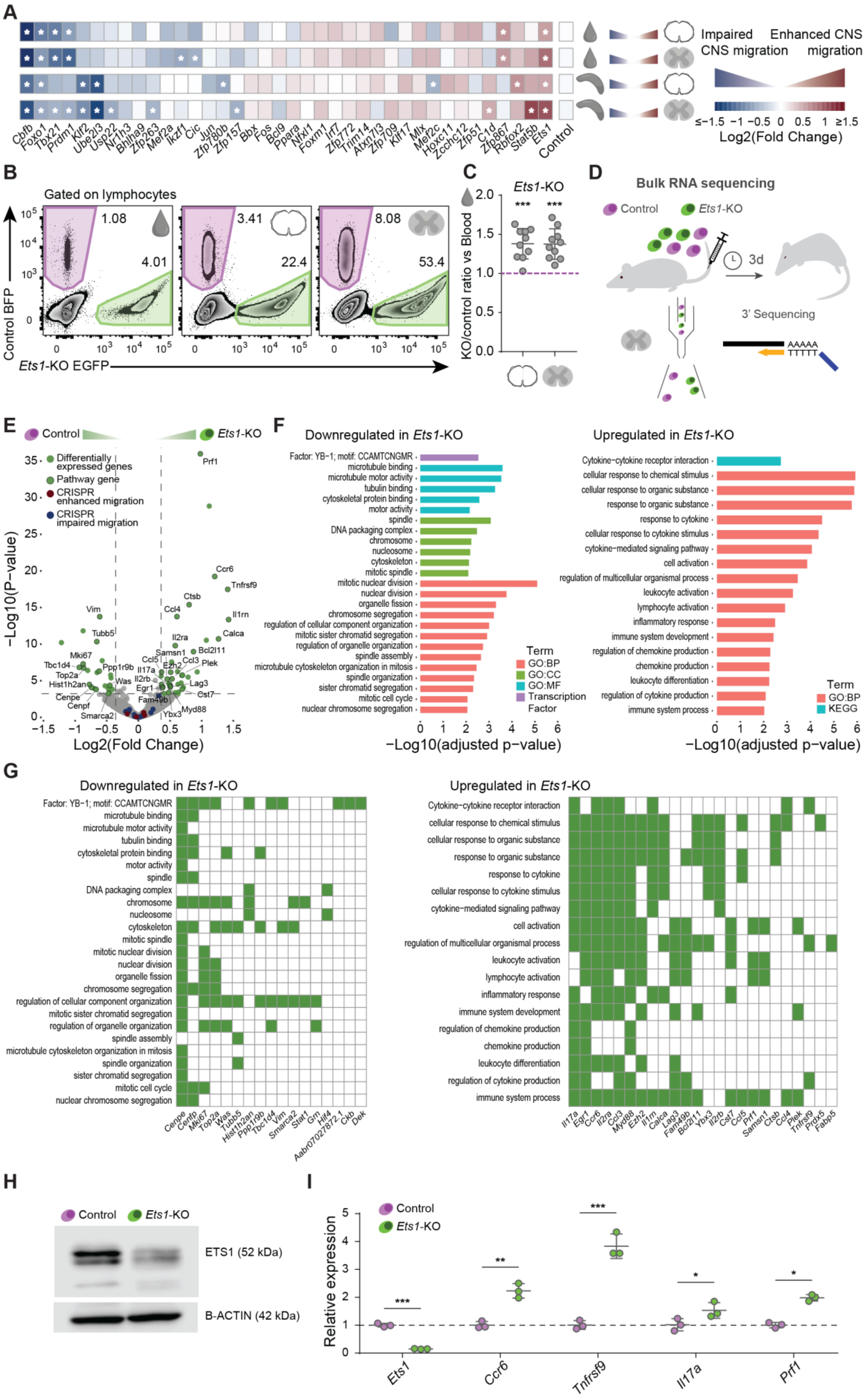
A
Impaired CNS migration
Enhanced CNS migration
Cbfb Foxo1 Tbx21 Prdm1 Klf2 Ube2l3 Usp22 Nr1h3 Bhlha9 Zfp263 Mef2a Ikzf1 Cic Jun Zfp780b Zfp157 Bbx Fos Bcl9 Ppara Nfxl1 Foxm1 Irf7 Zfp772 Trim14 Atxn7l3 Zfp709 Klf17 Mlx Mef2c Hoxc11 Zcchc12 Zfp51 C1d Zfp867 Rbfox2 Stat5b Ets1 Control
≤−1.5 −1 −0.5 0 0.5 1 ≥1.5
Log2(Fold Change)
B
Gated on lymphocytes
1.08
4.01
3.41
22.4
8.08
53.4
Control BFP
Ets1-KO EGFP
C
Ets1-KO
KO/control ratio vs Blood
D
Bulk RNA sequencing
Control
Ets1-KO
3d
3' Sequencing
AAAAA
TTTTT
E
Control
Ets1-KO
Differentially expressed genes
Pathway gene
CRISPR enhanced migration
CRISPR impaired migration
−Log10(P−value)
Log2(Fold Change)
F
Downregulated in Ets1-KO
Upregulated in Ets1-KO
Factor: YB−1; motif: CCAMTCNGMR
microtubule binding
microtubule motor activity
tubulin binding
cytoskeletal protein binding
motor activity
spindle
DNA packaging complex
chromosome
nucleosome
cytoskeleton
mitotic spindle
mitotic nuclear division
nuclear division
organelle fission
chromosome segregation
regulation of cellular component organization
mitotic sister chromatid segregation
regulation of organelle organization
spindle assembly
microtubule cytoskeleton organization in mitosis
spindle organization
sister chromatid segregation
mitotic cell cycle
nuclear chromosome segregation
Term
GO:BP
GO:CC
GO:MF
Transcription Factor
Cytokine−cytokine receptor interaction
cellular response to chemical stimulus
cellular response to organic substance
response to organic substance
response to cytokine
cellular response to cytokine stimulus
cytokine−mediated signaling pathway
cell activation
regulation of multicellular organismal process
leukocyte activation
lymphocyte activation
inflammatory response
immune system development
regulation of chemokine production
chemokine production
leukocyte differentiation
regulation of cytokine production
immune system process
Term
GO:BP
KEGG
−Log10(adjusted p−value)
G
Downregulated in Ets1-KO
Upregulated in Ets1-KO
Cenpe Cenpf Mki67 Top2a Was Tubb5 Hist1h2an Ppp1r9b Tbc1d4 Vim Smarca2 Stat1 Grn Hif4 Aabr07027872.1 Ckb Dek
Il17a Egr1 Ccr6 Il2ra Ccl3 Myd88 Ezh2 Il1rn Calca Lag3 Fam49b Bcl2l11 Ybx3 Il2rb Cst7 Ccl5 Prf1 Samsn1 Ctsb Ccl4 Plek Tnfrsf9 Prdx5 Fabp5
H
Control
Ets1-KO
ETS1 (52 kDa)
B-ACTIN (42 kDa)
I
Control
Ets1-KO
Relative expression
Ets1
Ccr6
Tnfrsf9
Il17a
Prf1

**Figure S3. ETS1 inhibits CD4$^+$ T cell migration to the CNS.**

**A,** Log2(Fold Change) heatmaps depicting the $T_{MBP}$ cell migratory phenotype of gene KOs as per the validation screen results for transcription factors (GO.0003700, GO.0003713 and GO.0003714). Only genes with a p-value < 0.01 and ≥ 3 "neg/pos|goodsgrna" as per the validation screen results are shown. **B,** Representative flow cytometry plots of $T_{MBP}$ cells recovered from blood, meninges and parenchyma after a co-transfer experiment with *Ets1*-KO and control cells. **C,** Migratory phenotype of *Ets1*-KO cells compared to control, shown as the ratio of *Ets1*-KO cell numbers to control cell numbers in meninges (left) or parenchyma (right) divided by the KO/control ratio in blood. A ratio of 1 indicates the migration behaviour of control $T_{MBP}$ cells, a ratio above 1 indicates enhanced migration into the CNS tissue. n = 10 rats. **D,** Experimental design of the bulk RNAseq experiment of *Ets1*-KO and control cells from the parenchyma of co-transferred animals. **E,** Volcano plot of the RNAseq results of *Ets1*-KO cells compared to control $T_{MBP}$ cells, lines indicate adjusted p-value = 0.05 and log2(Fold change) = ±3 standard deviations of the sample. Green, significantly differentially expressed genes; blue, essential genes "facilitating" CNS migration in $T_{MBP}$ cells ("CRISPR impaired migration"); red, essential genes "braking" CNS migration ("CRISPR enhanced migration"). Only significant genes belonging to pathways of the subsequent analysis are labelled. **F,** Pathway analysis by gProfiler of the top downregulated pathways (left panel) and top upregulated pathways (right panel) in the *Ets1*-KO compared to control $T_{MBP}$ cells. **G,** Significant (adjusted P < 0.05 and absolute log2(Fold change) > 3 standard deviations of the sample) genes belonging to the pathways in **f**. **H,** Western blot showing the downregulation of ETS1 protein levels in the KO cells. **I,** qPCR validation of selected upregulated genes in the *Ets1*-KO cells driving the upregulated pathways. n = 3 samples per group. **C,** One sample t-test against hypothetical mean = 1; **I**, multiple paired t-tests with two-stage linear step-up procedure of Benjamin, Krieger and Yekutieli (fdr 1%) multiple testing correction. Figures show mean ± s.d, P > 0.05 ns (non-significant), P < 0.05 *, P < 0.01 **, P < 0.001 ***, P < 0.0001 ****.

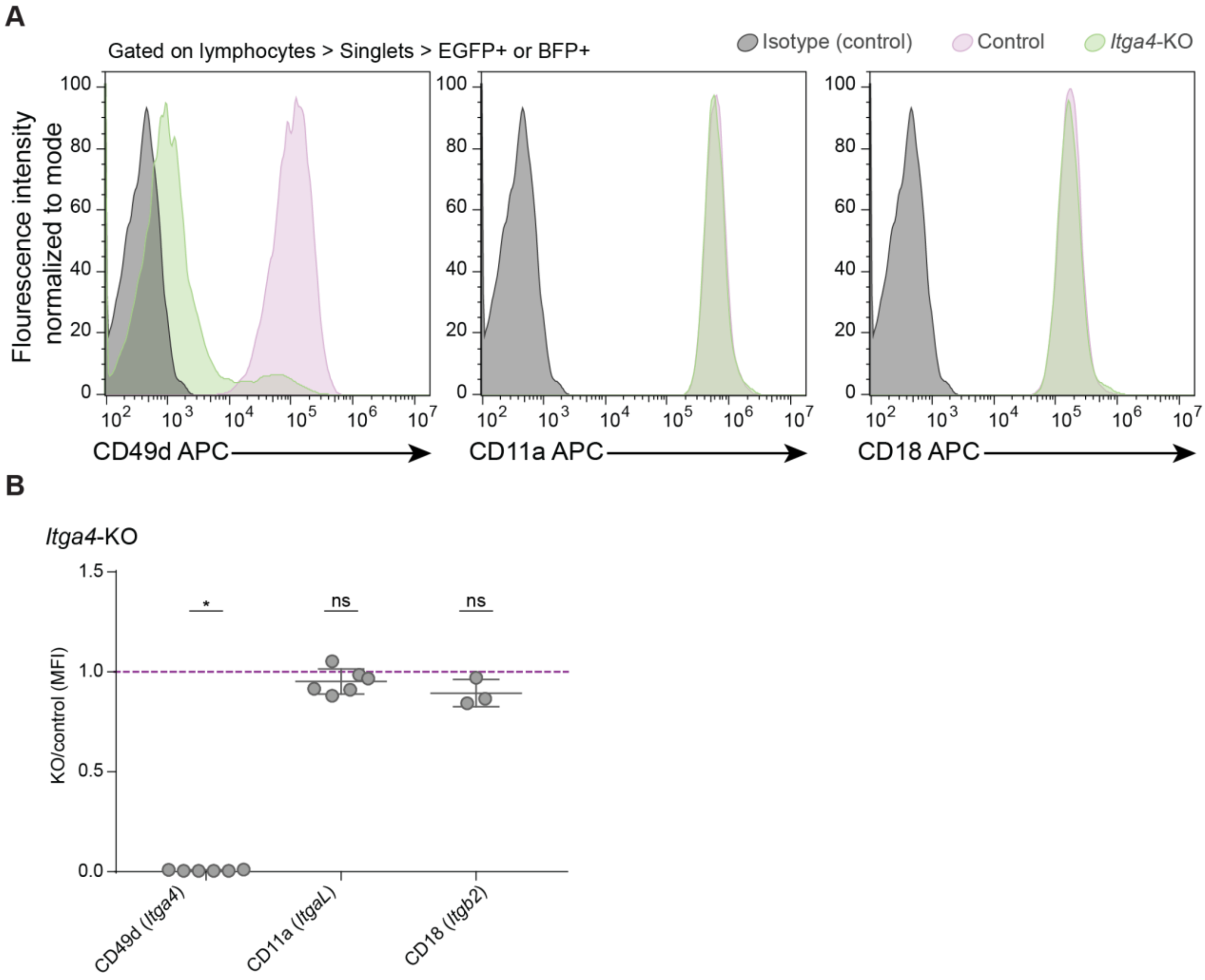


**Figure S4. *Itga4*-KO does not downregulate surface expression of integrins.**

**A,** Representative flow cytometry plots showing in grey the isotype-matched antibody labelling control, in lilac the control $T_{MBP}$ cells, and in green the *Itga4*-KO $T_{MBP}$ cells, for CD49d (*Itga4*), CD11a (*ItgaL*) and CD18 (*Itgb2*). B**,** Quantification of the integrin labelling median fluorescence intensity of the *Itga4*-KO cells, normalized to the control intensity. A ratio of 1 indicates the surface integrin expression levels of control $T_{MBP}$ cells, a ratio below 1 indicates reduced integrin surface expression. n = 6 (*Itga4*-KO CD49d and CD11a), n = 3 (*Itga4*-KO CD18), all independent stainings. **B**, One sample t-test or Wilcoxon signed-rank test against hypothetical mean = 1. Figures show mean ± s.d, P > 0.05 ns (non-significant), P < 0.05 *, P < 0.01 **, P < 0.001 ***, P < 0.0001 ****.

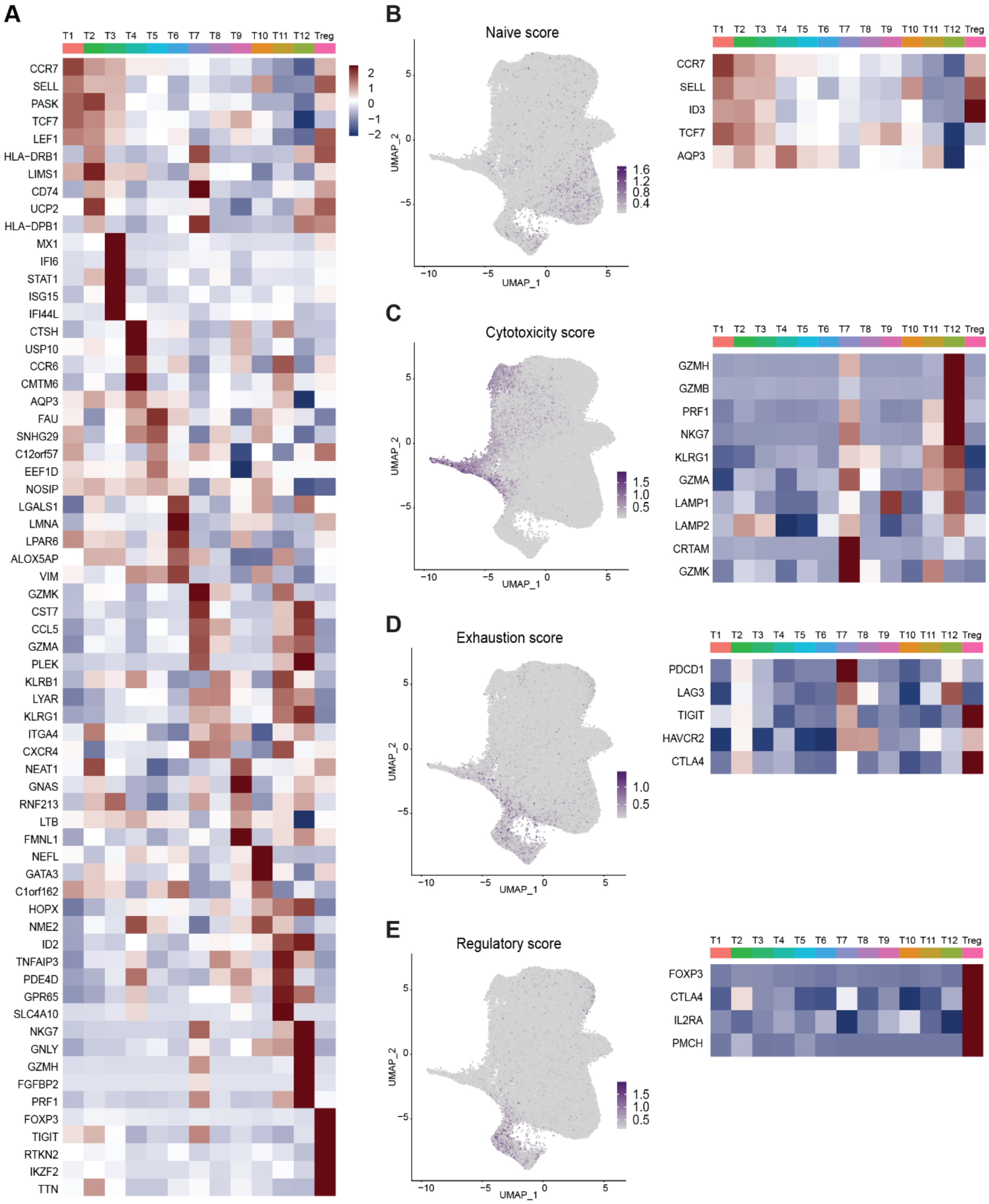


**Figure S5. Molecular characterization of CD4$^{+}$ T cell clusters**

**A,** Heatmap showing the relative expression levels of the top five differentially regulated genes per cluster. **B-E,** Naïve (**B**), cytotoxicity (**C**), exhaustion (**D**) and regulatory (**E**) T cell marker gene expression, across cells (left) and clusters (right).

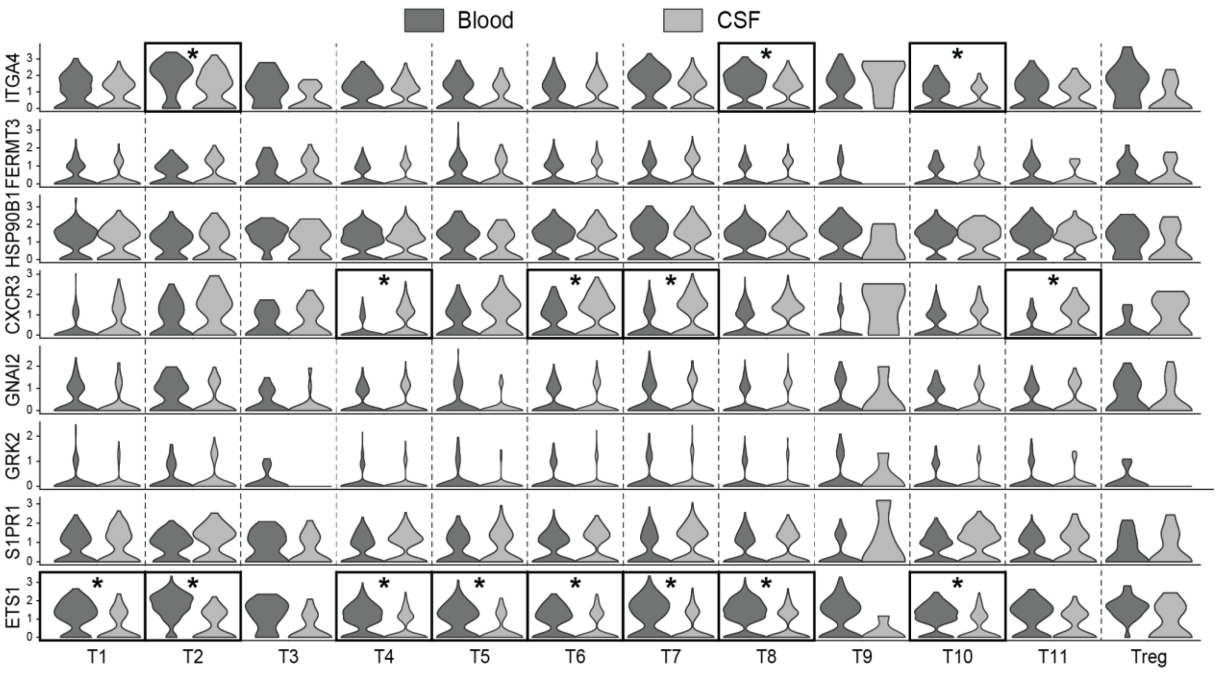


**Figure S6. Expression pattern of essential regulators in human CD4⁺ T cells from blood and CSF**

Violin plots comparing selected gene expression levels between T cells (analysed per cluster) from blood (dark gray) and CSF (light gray) in MS patients for those $CD4^+$ T cell clones whose TCR sequences were detected in blood and CSF. Stars indicate significance as per adjusted P < 0.05 and absolute log2(Fold Change) > 3 times the standard deviation of the sample.
